# Adaptation to plant shade relies on rebalancing the transcriptional activity of the PIF-HFR1 regulatory module

**DOI:** 10.1101/2020.09.02.271551

**Authors:** Sandi Paulišić, Christiane Then, Benjamin Alary, Fabien Nogue, Miltos Tsiantis, Jaime F. Martínez-García

## Abstract

Shade caused by the proximity of neighboring vegetation triggers a set of acclimation responses to either avoid or tolerate shade. Comparative analyses between the shade avoider *Arabidopsis thaliana* and the shade tolerant *Cardamine hirsuta*, revealed a role for the atypical basic-helix-loop-helix LONG HYPOCOTYL IN FR 1 (HFR1) in maintaining the shade-tolerance in *C. hirsuta*, inhibiting hypocotyl elongation in shade and constraining expression profile of shade induced genes. We showed that *C. hirsuta* HFR1 protein is more stable than its *A. thaliana* counterpart, contributing to enhance its biological activity. The enhanced HFR1 activity is accompanied by an attenuated PHYTOCHROME INTERACTING FACTOR (PIF) activity in *C. hirsuta*. As a result, the PIF-HFR1 module is imbalanced, causing a reduced PIF activity and attenuating other PIF-mediated responses such as warm temperature-induced hypocotyl elongation (thermomorphogenesis) and dark-induced senescence. By this mechanism and that of the already-known of phytochrome A photoreceptor, plants might ensure to properly adapt and thrive in habitats with disparate light amounts.

## INTRODUCTION

Acclimation of plants to adjust their development to the changing environment is of uttermost importance. This acclimation relies on the plant’s ability to perceive many cues such as water, nutrients, temperature or light. Conditions in nature often involve simultaneous changes in multiple light cues leading to an interplay of various photoreceptors to adjust plant growth appropriately ^1-5^. Nearby vegetation can impact both light quantity and quality. Under a canopy, light intensity is decreased and its quality is changed as the overtopping green leaves strongly absorb blue and red light (R) but reflect far-red light (FR). As a consequence, plants growing in forest understories receive less light of a much lower R to FR ratio (R:FR) than those growing in open spaces. In dense plant communities, FR reflected by neighboring plants also decreases R:FR but typically without changing light intensity. We refer to the first situation as canopy shade (very low R:FR) and the second as proximity shade (low R:FR). In general, two strategies have emerged to deal with shade: avoidance and tolerance ^4,6,7^. Shade avoiders usually promote elongation of organs to outgrow the neighbors and avoid light shortages, reduce the levels of photosynthetic pigments to cope to light shortage, and accelerate flowering to ensure species survival ^8^. The set of responses to acclimate to shade is collectively known as the shade avoidance syndrome (SAS). In contrast, shade-tolerant species usually lack the promotion of elongation growth in response to shade and have developed a variety of traits to acclimate to low light conditions and optimize net carbon gain ^7,9^.

In *Arabidopsis thaliana*, a shade avoider plant, low R:FR is perceived by phytochromes. Among them, phyA has a negative role in elongation, particularly under canopy shade, whereas phyB inhibits elongation inactivating PHYTOCHROME INTERACTING FACTORS (PIFs), members of the basic-helix-loop-helix (bHLH) transcription factor family that promote elongation growth. In particular, PIFs induce hypocotyl elongation by initiating an expression cascade of genes involved in auxin biosynthesis and signaling [e.g., *YUCCA 8* (*YUC8*), *YUC9, INDOLE-3-ACETIC ACID INDUCIBLE 19* (*IAA19*), *IAA29*] and other processes related to cell elongation [e.g., *XYLOGLUCAN ENDOTRANSGLYCOSYLASE 7* (*XTR7*)]. Genetic analyses indicated that PIF7 is the key PIF regulator of the low R:FR-induced hypocotyl elongation. Indeed, *pif7* mutant responds poorly to low R:FR compared to the *pif4 pif5* double or *pif1 pif3 pif4 pif5* quadruple (*pifq*) mutants ^10,11^. PhyB-mediated shade signaling involves other transcriptional regulators, such as *LONG HYPOCOTYL IN FR 1* (*HFR1*), *PHYTOCHROME RAPIDLY REGULATED 1* (*PAR1*), *BIM1, ATHB4* or *BBX* factors, that either promote or inhibit shade-induced hypocotyl elongation ^12-18^. HFR1, a member of the bHLH family, is structurally related to PIFs but lacks the phyB- and DNA-binding ability that PIFs possess ^19,20^. HFR1 inhibits PIF activity by heterodimerizing with them, as described for PIF1 ^21^, PIF3 ^22^, PIF4 and PIF5 ^23^, Heterodimerization with HFR1 prevents PIFs from binding to the DNA and altering gene expression. In this manner HFR1 acts as a transcriptional cofactor that modulates SAS responses, e.g. it inhibits hypocotyl elongation in seedlings in a PIF-dependent manner, forming the PIF-HFR1 transcriptional regulatory module ^19^.

What mechanistic and regulatory adjustments in shade signaling are made between species to adapt to plant shade is a topic that has not received much attention until now. This question has been recently addressed performing comparative analyses between phylogenetically related species. In two related *Geranium* species that showed petioles with divergent elongation responses to shade, transcriptomic analysis led to propose that differences in expression of three factors, *FERONIA, THESEUS1* and *KIDARI*, shown to activate SAS elongation responses in *A. thaliana*, might be part of the adjustments necessary to acquire a shade-avoiding or tolerant habit ^24^. When comparing two related mustard species that showed divergent hypocotyl elongation response to shade, *A. thaliana* and *Cardamine hirsuta* ^25^, molecular and genetic analyses indicated that phyA, and to a lesser extent phyB, contributed to establish this divergent response. In particular, the identification and characterization of the *C. hirsuta* phyA-deficient *slender in shade 1* (*sis1*) mutant indicated that differential features of this photoreceptor in *A. thaliana* and *C. hirsuta* could explain their differential response to shade. Thus, stronger phyA activity in *C. hirsuta* wild-type plants resulted in a suppressed hypocotyl elongation response when exposed to low or very low R:FR ^26^. These approaches indicated that the implementation of shade avoidance and shade tolerance involved the participation of shared genetic components.

With this frame of reference, we asked whether the phyB-dependent PIF-HFR1 module was also relevant to shape the shade response habits in different plant species. We found that *C. hirsuta* plants deficient in ChHFR1 gained a capacity to elongate in response to shade. We also report that *AtHFR1* and *ChHFR1* are expressed at different levels and encode proteins with different protein stability. We propose that adaptation to plant shade in *A. thaliana* and *C. hirsuta* relies on the PIF-HFR1 regulatory module.

## RESULTS

### *HFR1* is required for the shade tolerance habit of *C. hirsuta*

First, we wanted to determine if HFR1 has a role in the shade-tolerance habit of *C. hirsuta*, i.e., whether *ChHFR1* contributes to inhibit hypocotyl elongation when this species is exposed to shade. For this purpose, we generated several *C. hirsuta* RNAi lines to downregulate *HFR1* expression (RNAi-HFR1 lines). As expected, *ChHFR1* expression was attenuated in seedlings of two RNAi-HFR1 selected lines (#01 and #21) compared to the wild type (Ch^WT^) (**Supplemental Figure 1a**). When growing under white light (W) of high R:FR (>1.5), hypocotyl length of these two RNAi-HFR1 lines was undistinguishable from Ch^WT^ (**Figure 1a)**. By contrast, under W supplemented with increasing amounts of FR (W+FR) resulting in moderate (0.09), low (0.05-0.06) and very low (0.02) R:FR (that simulated proximity and canopy shade), the hypocotyl elongation of RNAi-HFR1 seedlings was significantly promoted compared to Ch^WT^, which was unresponsive (**Figure 1a**).

**Figure 1.**
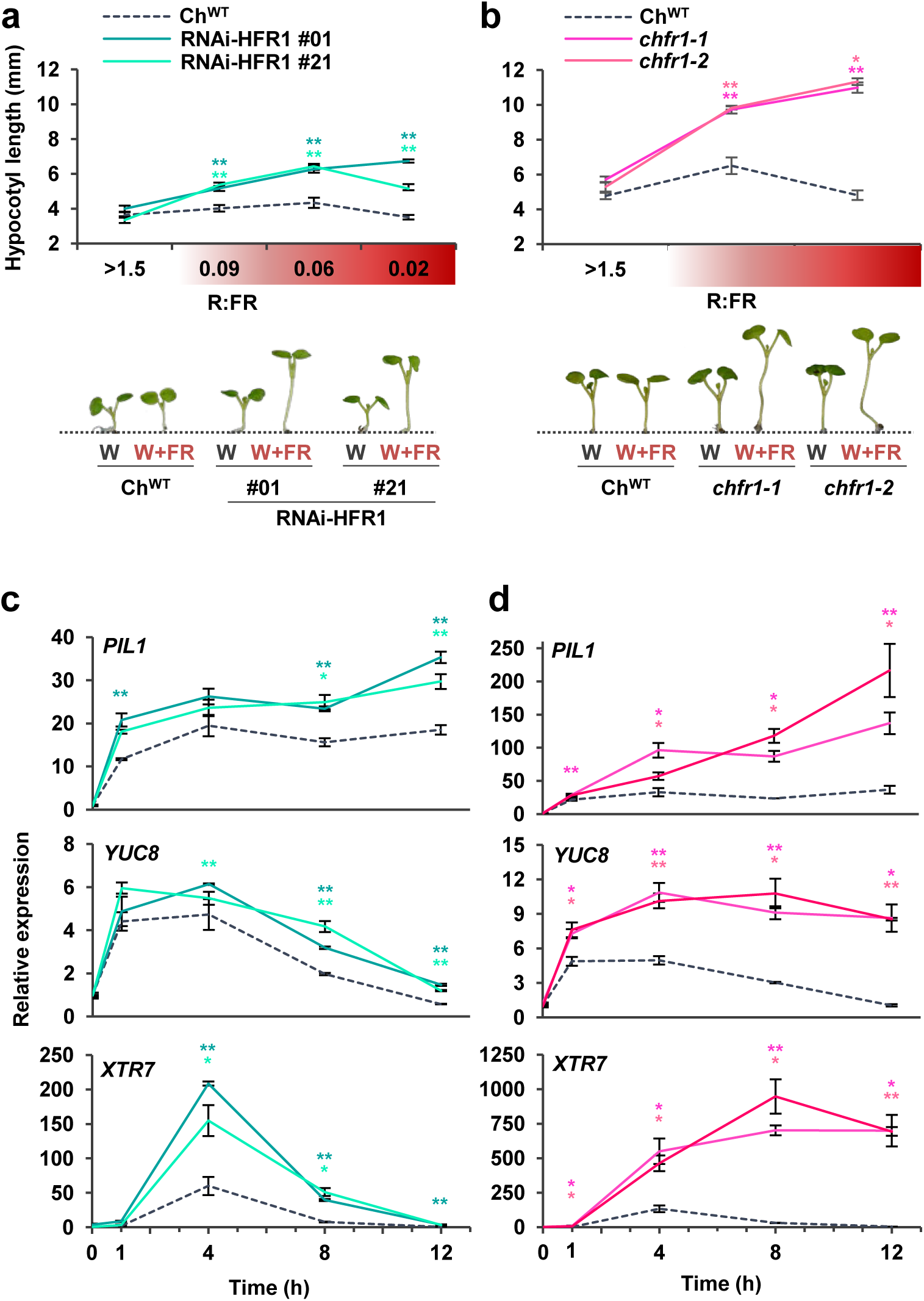
Hypocotyls of *C. hirsuta* seedlings with reduced levels of *ChHFR1* strongly elongate in response to simulated shade. Hypocotyl length of Ch^WT^, **(a)** RNAi-ChHFR1 transgenic and **(b)** *chfr1* mutant seedlings grown under different R:FR. Seedlings were grown for 7 days in continuous W (R:FR>1.5) or for 3 days in W then transferred to W supplemented with increasing amounts of FR (W+FR) for 4 more days, producing various R:FR. Aspect of representative 7-day old Ch^WT^, RNAi-HFR1 and *chfr1-1* seedlings grown in W or W+FR (R:FR, 0.02), as indicated, is shown in lower panel. Effect of W+FR exposure on the expression of *PIL1, YUC8* and *XTR7* genes in seedlings of Ch^WT^, **(c)** RNAi-HFR1 and **(d)** c*hfr1* mutant lines. Expression was analyzed in 7-day old W-grown seedlings transferred to W+FR (R:FR, 0.02) for 0, 1, 4, 8 and 12 h. Transcript abundance is normalized to *EF1α* levels. Values are the means ± SE of three independent biological replicates relative to Ch^WT^ value at 0 h. Asterisks mark significant differences (Student *t*-test: ** p-value <0.01; * p-value <0.05) relative to Ch^WT^ value at the same time point.

Using CRISPR-Cas9, we obtained two mutant lines of *ChHFR1* (named *chfr1-1* and *chfr1-2*) with a single nucleotide insertion in their sequence leading to a premature stop codon (**Supplemental Figure 1c**). These mutants showed a non-significant decrease of *ChHFR1* expression in W-grown seedlings (**Supplemental Figure 1b**). Similar to the RNAi-HFR1 lines, their hypocotyls were undistinguishable from Ch^WT^ under W but elongated strongly in response to W+FR exposure (**Figure 1b)**, showing a *slender in shade* (*sis*) phenotype. Together, we concluded that HFR1 represses hypocotyl elongation in response to shade in *C. hirsuta*.

Exposure of *A. thaliana* wild-type (At^WT^) and Ch^WT^ seedlings to low R:FR induces a rapid increase in the expression of various direct target genes of PIFs, including *PIF3-LIKE 1* (*PIL1*), *YUC8* and *XTR7* (**Figure 1c, d**) ^26-28^. The shade-induced expression of these genes was significantly higher in RNAi-HFR1 and *chfr1* mutant lines compared to Ch^WT^ (**Figure 1c, d**), indicating that ChHFR1 might repress shade-triggered hypocotyl elongation in part by down-regulating the rapid shade-induced expression of these genes in *C. hirsuta*, as it was observed with AtHFR1 in *A. thaliana* seedlings ^23^.

### *HFR1* expression is higher in *C. hirsuta* than in *A. thaliana* seedlings

To test if the lack of elongation of Ch^WT^ hypocotyls in response to shade was caused by higher levels of *ChHFR1* expression in this species, we used primer pairs that amplify *HFR1* (**Supplemental Figure 2a**) and three housekeeping genes (*EF1α, SPC25, YLS8*) in both species ^26^. As expected, expression of *HFR1* was induced in shade-treated seedlings of both species, in agreement with the presence of canonical PIF-binding sites (G-box, CACGTG) in the *HFR1* promoters ^23,29^ (**Supplemental Figure 3a**). More importantly, *ChHFR1* transcript levels were always higher than those of *AtHFR1* during the whole period analyzed (from days 3 to 7) (**Figure 2**). Because HFR1 is part of the PIF-HFR1 regulatory module, we next compared transcript levels of *PIF* genes in both species. *PIF7* expression was significantly lower in *C. hirsuta* than in *A. thaliana* in either W or W+FR during the period analyzed (**Figure 2**). By contrast, *PIF4* expression was higher in *C. hirsuta* than in *A. thaliana*, whereas that of *PIF5* was similar in both species (**Supplemental Figure 2b**). Together, these results indicated that whereas *HFR1* expression is enhanced, that of *PIF7* is globally attenuated in Ch^WT^ compared to At^WT^ seedlings. As a consequence, the PIF-HFR1 transcriptional module might be differently balanced in these species, with HFR1 imposing a stronger suppression on the PIF7-driven hypocotyl elongation in the shade-tolerant *C. hirsuta* seedlings.

**Figure 2.**
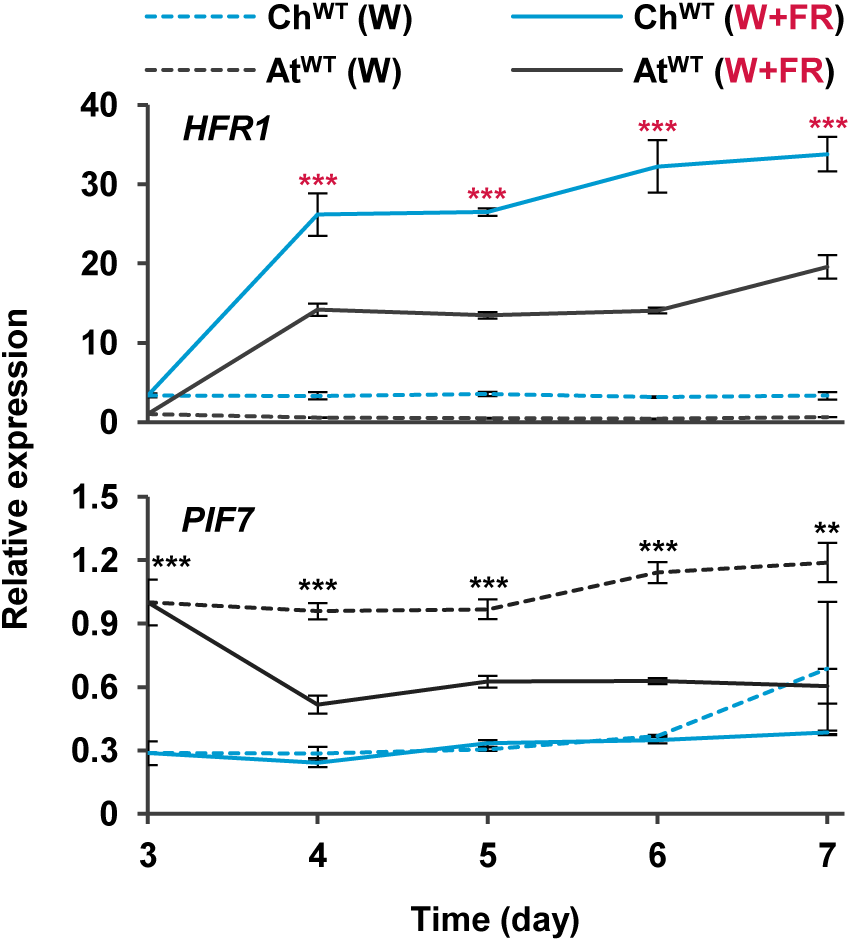
Levels of *HFR1* transcript are higher in *C. hirsuta* than *A. thaliana* seedlings. Seedlings of Ch^WT^ and At^WT^ were grown for 3 days in W then either kept under the same conditions or transferred to W+FR (R:FR, 0.02) for the indicated times. Plant material was harvested every 24 h. Transcript abundance of *HFR1* and *PIF7* was normalized to three reference genes (*EF1α, SPC25*, and *YLS8*). Expression values are the means ± SE of three independent biological replicates relative to the data of At^WT^ grown in continuous W at day 3. Asterisks mark significant differences (2-way ANOVA: * p-value <0.05, ** p-value <0.01, *** p-value <0.001) between Ch^WT^ and At^WT^ when grown under W (black asterisks) or W+FR (red asterisks).

### ChHFR1 protein is more stable than AtHFR1

A higher intrinsic activity of ChHFR1 compared to its orthologue AtHFR1 might also contribute to the role of this transcriptional cofactor in maintaining the shade tolerance habit of *C. hirsuta*. To test this possibility, we transformed *A. thaliana hfr1-5* plants with constructs to express either *AtHFR1* or *ChHFR1* fused to the 3x hemagglutinin tag (3xHA). These genes were expressed under the transcriptional control of the 2 kb of the *AtHFR1* promoter (*pAt*), generating *hfr1>pAt:ChHFR1* and *hfr1>pAt:AtHFR1* lines (**Figure 3a**). Fusion of *pAt* to the *GUS* reporter gene resulted in GUS activity in cotyledons and roots of transgenic lines, with increased levels in hypocotyls of seedlings exposed for 2-4 h to W+FR (**Supplemental Figure 3b**). Several independent transgenic lines of each construct were analyzed for hypocotyl length, *HFR1* transcript levels and 3xHA-tagged protein abundance. In these lines, HFR1 biological activity was estimated as the difference in hypocotyl length of seedlings grown under W+FR (Hyp^W+FR^) and W (Hyp^W^) (Hyp^W+FR^-Hyp^W^) ^26^. The potential to suppress the hypocotyl elongation in shade below that of *hfr1-5* seedlings would depend on the transcript level of *HFR1* and/or its protein levels. The *hfr1>pAt:ChHFR1* lines had shorter hypocotyls in shade compared to *hfr1>pAt:AtHFR1* lines of similar *HFR1* expression levels (**Figure 3b, c**). Consistently with this, we observed much higher abundance of HFR1-3xHA protein after shade exposure in *hfr1>pAt:ChHFR1* lines than in *hfr1>pAt:AtHFR1* lines with comparable levels of *HFR1* expression (**Figure 3d**), suggesting that the ChHFR1 protein might be much more stable.

**Figure 3.**
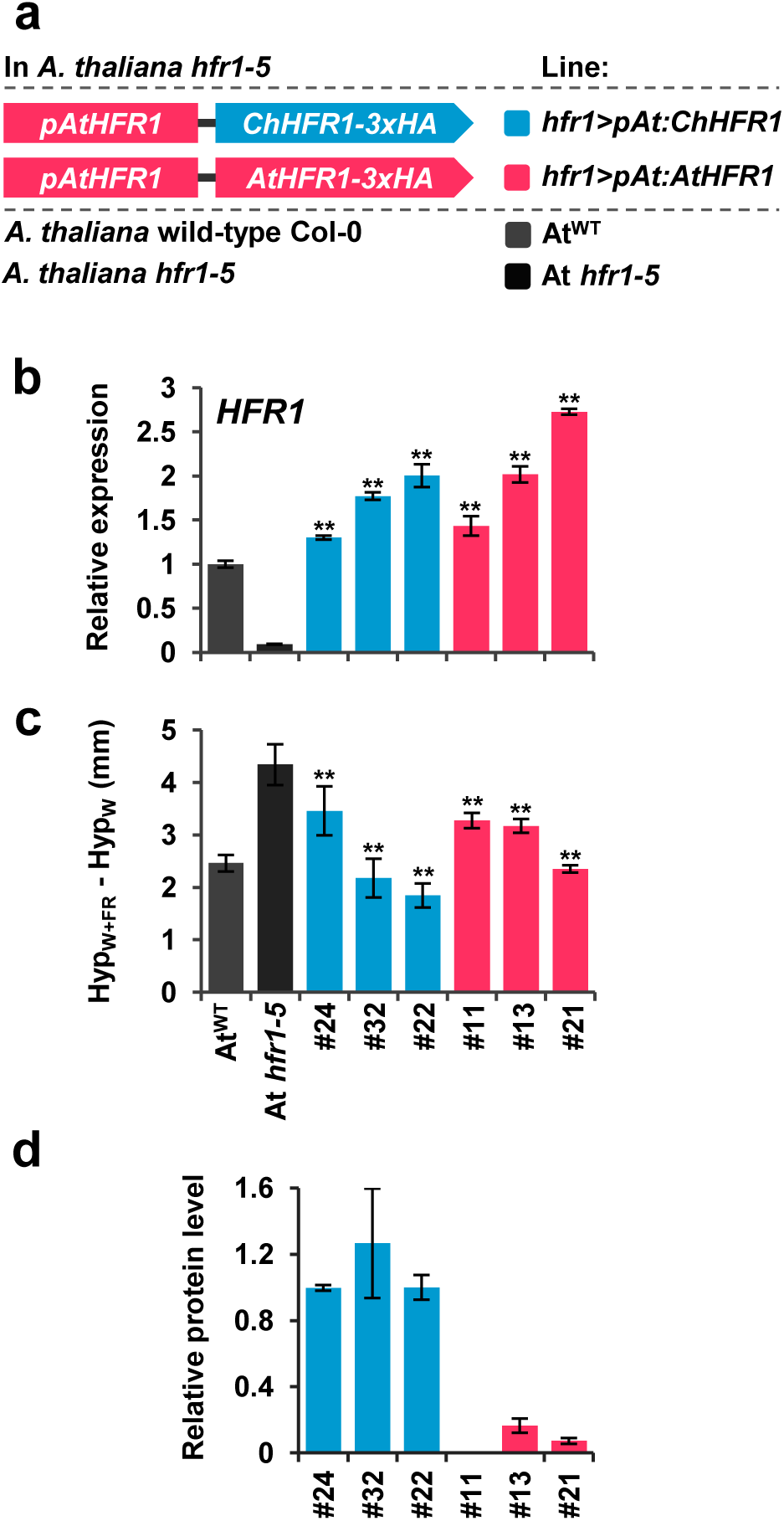
The activity of ChHFR1 is higher than that of AtHFR1 in *A. thaliana* seedlings. **(a)** Cartoon of constructs containing *ChHFR1* or *AtHFR1* under the *HFR1* promoter of *A. thaliana* (*pAtHFR1*) used to complement *hfr1-5* mutant of *A. thaliana* (At *hfr1-5*). **(b)** Relative expression of *HFR1* in seedlings of At^WT^, At *hfr1-*5, hfr1*>pAt:ChHFR1* (in blue) and *hfr1>pAt:AtHFR1* (in red) lines grown under W+FR (R:FR, 0.02). Expression values are the means ± SE of three independent biological replicates relative to the data of 7 days old At^WT^. Transcript abundance is normalized to *UBQ10* levels. **(c)** Elongation response of seedlings of the indicated lines grown for 7 days in continuous W or 2 days in W then transferred for 5 days to W+FR (R:FR, 0.02). The mean hypocotyl length in W (Hyp^W^) and W+FR (Hyp^W+FR^) of at least four biological replicates was used to calculate Hyp^W+FR^-Hyp^W^. **(d)** Relative HFR1 protein levels in seedlings of the indicated lines, normalized to actin protein levels, are the means ± SE of three independent biological replicates relative to *hfr1>pAt:ChHFR1* line #22 that is taken as 1. Seedlings were grown for 7 days in continuous W (∼20 µmol m^-2^ s^-1^) after which they were incubated for 3 h in high W (∼100 µmol m^-2^ s^-1^) and transferred to W+FR (R:FR, 0.06) for 3 h. Asterisks mark significant differences (Student *t*-test: ** p-value <0.01; * p-value <0.05) relative to At *hfr1-5*.

AtHFR1 stability is dependent on light conditions. In etiolated seedlings, exposure to W promotes stabilization and accumulation of AtHFR1, whereas in W-grown seedlings, high intensity of W increases its abundance ^30,31^. Importantly, AtHFR1 stability has a strong impact on its biological activity as overexpression of stable forms of this protein leads to phenotypes resulting from enhanced HFR1 activity ^19,31^. As AtHFR1 and ChHFR1 primary structures are globally similar (**Supplemental Figure 4a**), we aimed to test if ChHFR1 stability is also light-dependent. We first examined ChHFR1 protein accumulation in response to different W intensities in seedlings of an *A. thaliana hfr1-5* line that constitutively express *ChHFR1* (*hfr1>35S:ChHFR1*) (**Supplemental Figure 4b**). When grown in our normal W conditions (∼20 µmol m^-2^ s^-1^), these seedlings accumulated low but detectable levels of ChHFR1; when transferred to higher W intensity (∼100 µmol m^-2^ s^-1^), ChHFR1 levels increased 10-fold (**Supplemental Figure 4c**). As *ChHFR1* is expressed under the constitutive 35S promoter, these results indicate that ChHFR1 protein accumulation is induced by high W intensity, as it has been described for AtHFR1 ^31^. This prompted us to pretreat W-grown seedlings with 3 h of high W intensity in all our subsequent experiments to analyze ChHFR1 levels.

Next, we exposed *hfr1>pAt:ChHFR1* (line #22) and *hfr1>pAt:AtHFR1* (line #13) seedlings to W+FR (**Figure 4a**). Although *HFR1* expression in both lines was similarly induced after 3 h of W+FR, *hfr1>pAt:ChHFR1* line displayed higher levels of recombinant HFR1 protein compared to *hfr1>pAt:AtHFR1* line after 3-6 h of W+FR exposure (**Figure 4a**), suggesting a higher stability of the *C. hirsuta* protein compared to the *A. thaliana* orthologue. ChHFR1 protein is more abundant than AtHFR1 also when transiently expressed to comparable levels in *Nicotiana benthamiana* leaves (**Figure 4b, c**). This indicates that the higher abundance of ChHFR1 is an intrinsic property of the protein.

**Figure 4.**
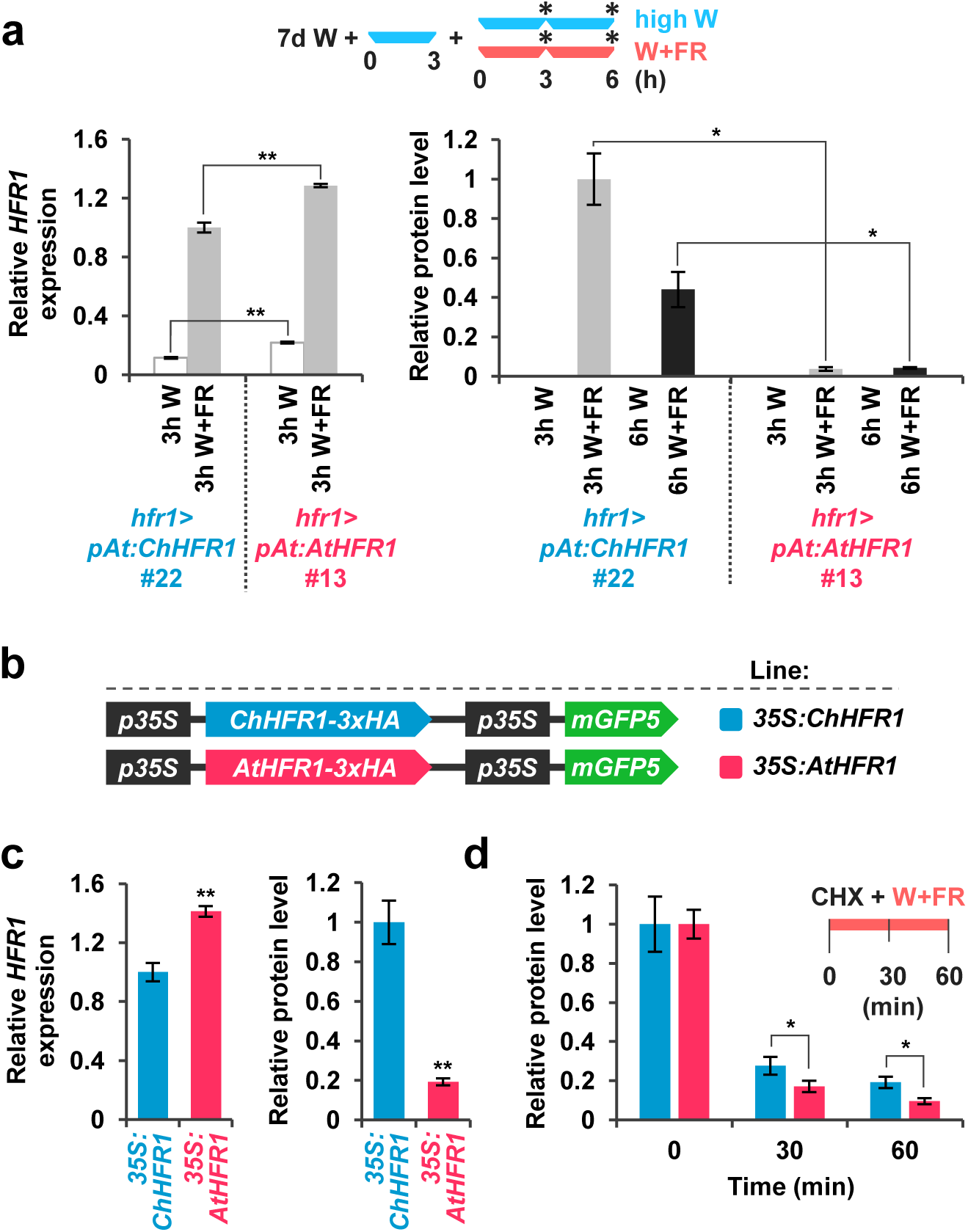
ChHFR1 and AtHFR1 proteins show different stability in shade. **(a)** Expression of *HFR1* and protein levels of HFR1-3xHA in seedlings of *hfr1>pAt:ChHFR1* (line #22) and *hfr1>pAt:AtHFR1* (line #13). Seedlings were grown for 7 days in continuous W (∼20 µmol m^-2^ s^-1^) after which they were incubated for 3 h in high W (∼100 µmol m^-2^ s^-1^) and then either kept at high W or transferred to W+FR (R:FR, 0.06) for 3 or 6 h, as indicated in the cartoon at the top. Relative *HFR1* transcript levels, normalized to *UBQ10*, are the means ± SE of three independent biological replicates relative to *hfr1>pAt:ChHFR1* #22 grown for 3 h under W+FR. Relative protein levels, normalized to actin, are the means ± SE of three independent biological replicates relative to *hfr1>pAt:ChHFR1* #22. Samples were collected at data points marked in the cartoon with asterisks. **(b)** Cartoon of constructs containing *ChHFR1* or *AtHFR1* under the *35S* promoter used for transient expression of transgenes in *N. benthamiana* leaves. **(c)** Relative *HFR1* transcript levels transiently expressed in tobacco leaves, normalized to the *GFP*, are the means ± SE of three independent biological replicates. Relative HFR1 protein levels, normalized to the GFP levels, are the means ± SE of four independent biological replicates. **(d)** Degradation of ChHFR1 (*35S:ChHFR1*) and AtHFR1 (*35S:AtHFR1*) in tobacco leaf discs treated with cycloheximide (CHX, 100 µM) for the indicated times. Tobacco plants were kept under high W (∼200 µmol m^-2^ s^-1^) for 3 days after agroinfiltration and then leaf circles were treated with W+FR (R:FR, 0.2) and CHX. Relative HFR1 protein levels, normalized to the GFP levels, are the means ± SE of four biological replicates relative to data point 0, taken as 1 for each line. Asterisks mark significant differences (2-way ANOVA: * p-value <0.05) between ChHFR1 and AtHFR1 at the same time point.

An increased ChHFR1 protein stability might be due to differences in degradation kinetics by the 26S proteasome. We addressed this possibility by treating tobacco leaf discs overexpressing *ChHFR1* and *AtHFR1* with the protein synthesis inhibitor cycloheximide (CHX) combined with shade (**Figure 4d**). This treatment resulted in a decrease in ChHFR1 and AtHFR1 protein levels. However, ChHFR1 degradation was significantly slower than that of AtHFR1 (**Figure 4d**), supporting that changes in degradation kinetics likely contribute to the observed differences in stability between ChHFR1 and AtHFR1.

### HFR1 interacts with PIF7

AtHFR1 has been shown to interact with all the members of the photolabile AtPIF quartet (PIF1, PIF3, PIF4 and PIF5). Using a yeast two-hybrid (Y2H) assay, we observed that AtHFR1 homodimerized, which indicated that its HLH domain is functional in this assay (**Figure 5a**). In the same assay, AtHFR1 was also shown to interact with AtPIF7 (**Figure 5a**). These results agree with recent data ^32^. Because AtPIF7 is the main PIF in *A. thaliana* promoting hypocotyl elongation in response to shade ^10^, we aimed to address whether HFR1 also interacts genetically with PIF7. First, we analyzed the genetic interaction between AtHFR1 and AtPIF7. After crossing *A. thaliana hfr1-5* with *pif7-1* and *pif7-2* mutants, we analyzed the hypocotyl response of the obtained double mutants in different low R:FR conditions. As expected, *hfr1* hypocotyls were longer and those of *pif7* mutants were shorter compared to At^WT^ under both W+FR conditions used (**Figure 5b**). In W and low R:FR (0.06), double *pif7 hfr1* mutant seedlings behaved mostly as *pif7* single mutants. However, under very low R:FR (0.02), they elongated similar to At^WT^ hypocotyls (**Figure 5b**). Together, these results indicate that *pif7* is epistatic over *hfr1* under proximity shade, whereas it seems more additive under canopy shade.

**Figure 5.**
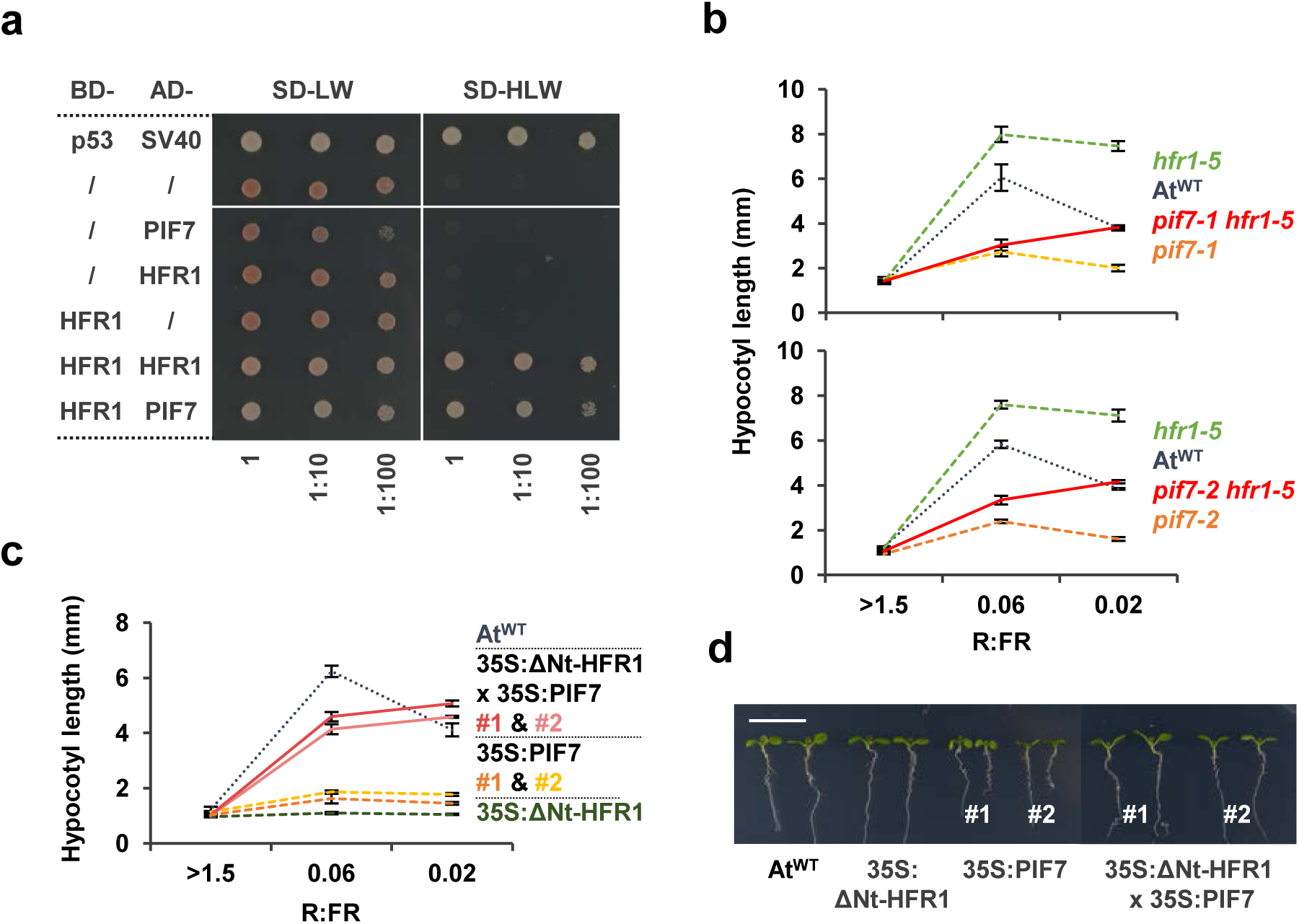
AtHFR1 interacts with AtPIF7. **(a)** Y2H growth assay showing the interaction between AtHFR1 and AtPIF7. The BD- and the AD-derivative constructs used in the assay are shown on the left side of the panel. SD-LW or SD-HLW refer to the selective medium (plated as drops in dilutions of 1, 1:10 and 1:100) indicative of transformed cells or interaction between the hybrid proteins, respectively. Truncated forms of murine p53 (BD-fused) and SV40 large T-antigen (AD-fused), known to interact, were used as a positive control. Empty vectors (/) were used as negative controls. Hypocotyl length of seedlings of At^WT^, **(b)** *pif7-1, hfr1-5, pif7-1 hfr1-5* (top graph), *pif7-2, hfr1-5* and *pif7-2 hfr1-5* (bottom graph) mutants, and **(c)** transgenic 35S:GFP-ΔNt-HFR1 (35S:ΔNt-HFR1), two lines of 35S:PIF7-CFP (35S:PIF7 #1 and #2), and 35S:GFP-ΔNt-HFR1 35S:PIF7-CFP double transgenic (35S:ΔNt-HFR1 × 35S:PIF7 #1 and #2) seedlings grown under different R:FR. Seedlings were grown in W (R:FR > 1.5) for 7 days or for 2 days in W and then transferred to two W+FR treatments (R:FR 0.06 or 0.02) for 5 additional days. Values of hypocotyl length are the means ± SE of three independent biological replicates (at least 10 seedlings per replica). **(d)** Aspect of representative 7-day-old W-grown seedlings shown in **c**.

To further analyze the HFR1-PIF7 interaction, we aimed to test if *HFR1* overexpression will interfere with *PIF7* overexpression and impede its effects. For HFR1, we used a line overexpressing a stable but truncated form of the protein (missing the N-terminal, 35S:GFP-ΔNt-HFR1, line #03) that strongly inhibits shade-induced hypocotyl elongation in *A. thaliana* without affecting other aspects of the seedling development ^19^ (**Figure 5c, d**). For PIF7 we used two available 35S:PIF7-CFP lines (#1 and #2) ^33^ that were almost unresponsive to W+FR (**Figure 5c**) and smaller and less developed than the At^WT^ in W (**Figure 5d**). In W, 35S:GFP-ΔNt-HFR1 35S:PIF7-CFP double transgenic seedlings (#1 and #2) did not differ in hypocotyl length and general aspect with At^WT^; interestingly they did elongate clearly in low and very low R:FR (**Figure 5c, d**). The recovery of the shade-induced hypocotyl elongation and size of the seedlings took place even though *HFR1* transcript levels were significantly lower than in the 35S:GFP-ΔNt-HFR1 parental line. *PIF7* transcript levels were not significantly different in the double transgenic seedlings than in their respective parental lines (**Supplemental Figure 5**). Therefore, the inhibitory effect of *PIF7-CFP* overexpression appeared to be counteracted by the overexpression of the truncated *HFR1*, further supporting the genetic interaction between HFR1 and PIF7 (**Figure 5c, d**).

Altogether, these analyses support that HFR1 and PIF7 interaction is important for the regulation of hypocotyl elongation in response to shade. These results are consistent with HFR1 functioning as a suppressor of PIF7.

### HFR1 restrains PIF activity in *C. hirsuta*

The similarity between shade-induced and warm temperature-induced hypocotyl elongation (thermomorphogenesis) suggests common underlying mechanisms. In *A. thaliana*, the increased activity of HFR1 at warm temperatures was previously shown to provide an important restraint on PIF4 action that drives elongation growth ^34^. Similarly, we hypothesized that the increased activity of HFR1 in *C. hirsuta* might restrain PIF activity more efficiently and consequently alter thermomorphogenesis (**Figure 6a**). We analyzed this response by growing seedlings constantly at 22°C, 28°C, or transferred from 22°C to 28°C after day 2 (**Figure 6b**). Whereas warm temperature promoted hypocotyl elongation of At^WT^ seedlings compared to those growing at 22°C, *pifq* and *pif7-2* mutant seedlings were almost unresponsive to 28°C, in accordance with the role of PIF4, PIF5 and PIF7 in thermomorphogenesis ^35-37^. Unlike the *hfr1-5* mutant, which was slightly but significantly more responsive than At^WT^, *A. thaliana* seedlings that overexpress a stable form of HFR1 (35S:GFP-ΔNt-HFR1, ΔNtHFR1) were almost unresponsive to 28°C (**Figure 6c**), indicating that HFR1 activity impacts this PIF-dependent response. A lack of hypocotyl elongation was also observed in Ch^WT^ at 28°C, a response that was recovered in the *C. hirsuta chfr1* mutant seedlings (**Figure 6c**). These results support our hypothesis that a strong suppression of PIFs by the enhanced HFR1 activity is responsible for the lack of hypocotyl elongation at 28°C of Ch^WT^ seedlings (**Figure 6a**). Together, our results suggest that the activity of the PIF-HFR1 regulatory module might be a general mechanism to coordinate the hypocotyl elongation in response to both W+FR exposure and 28°C.

**Figure 6.**
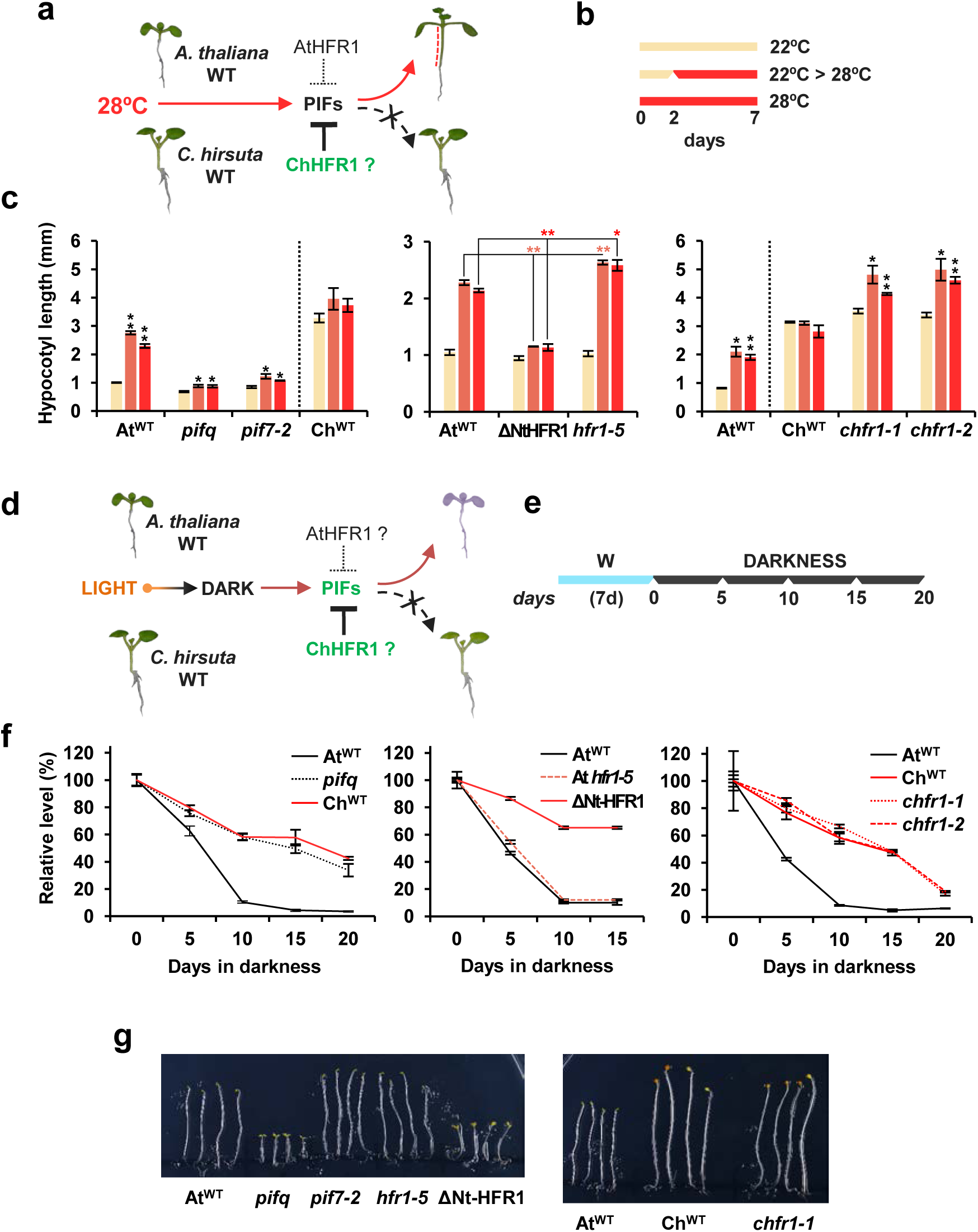
*C. hirsuta* has an attenuated hypocotyl elongation at warm temperature and delayed dark-induced senescence (DIS). **(a)** In At^WT^, PIFs promote hypocotyl elongation as a response to warm temperature (28°C). High ChHFR1 activity is expected to inhibit this response by repressing PIFs more effectively in Ch^WT^ and attenuate hypocotyl elongation at 28°C. **(b)** Seedlings were grown for 7 days in W at either 22°C, 2 days at 22°C then transferred to 28°C for additional 5 days (22°C > 28°C) or for 7 days at 28°C, as represented in the panel. **(c)** Hypocotyl length of seedlings of (left) At^WT^, *pifq, pif7-2*, Ch^WT^, (middle) 35S:GFP-ΔNt-HFR1 (ΔNtHFR1), *hfr1-5* and (right) *chfr1-1* and *chfr1-2* lines grown at warm temperatures. **(d)** In At^WT^, PIF-mediated DIS involves a reduction of chlorophyll levels. HFR1 activity might inhibit DIS through repression of PIFs. If PIF activity is attenuated in Ch^WT^, DIS would be delayed in this species compared to At^WT^. **(e)** Seedlings were grown for 7 days in W and then transferred to total darkness for several days to induce senescence, as illustrated at the right panel. **(f)** Relative chlorophylls levels of (left) At^WT^, *pifq*, Ch^WT^, (middle) ΔNtHFR1, *hfr1-5* and (right) *chfr1-1* and *chfr1-2* lines after DIS was promoted for the indicated time. For each genotype, data are relative to pigment levels at time 0 (7 days in W). **(g)** Aspect of 4-day old dark-grown seedlings of At^WT^, *pifq, pif7-2, hfr1-5* and ΔNt-HFR1 (left panel), and At^WT^, Ch^WT^ and *chfr1-1* (right panel).

We also studied dark-induced senescence (DIS), another PIF-dependent process (**Figure 6d**). In *A. thaliana*, DIS can be induced by transferring light grown seedlings to complete darkness, a process in which PIF4 and PIF5 have major roles ^38-40^. DIS results in a degradation of chlorophylls, which can be quantified as markers of senescence progression ^38,40^. To examine DIS, we transferred light-grown At^WT^, *pifq* and Ch^WT^ seedlings to total darkness for up to 20 days (**Figure 6e**). After DIS was activated, At^WT^ seedlings became pale and eventually died. After just 5 days of darkness, chlorophyll levels dropped, and longer dark treatments resulted in pronounced differences between the three genotypes. At^WT^ seedlings became visibly yellow at day 10, accompanied by a strong reduction of chlorophyll levels that dropped to less than 10% (**Figure 6f**). DIS was delayed in 35S:GFP-ΔNt-HFR1 seedlings, supporting that a stable HFR1 form can interfere with PIF activity in regulating this trait. However, DIS in was not advanced in *hfr1* mutants (**Figure 6e**). In Ch^WT^ seedlings, chlorophyll levels declined more slowly and seedlings were still green after 20 days of darkness, just like *pifq* (**Figure 6e**). The observed delay in the DIS in *C. hirsuta* was not affected in *chfr1* mutants, suggesting that HFR1 does not regulate this trait in any of the two species. It also pointed to a reduced PIF activity as the main cause for the delayed DIS in this species (**Figure 6d-f**). As HFR1 is very unstable, particularly in dark-grown conditions ^30,31^, it seems plausible that HFR1 does not accumulate in seedlings when transferred to the dark. Despite this attenuation of PIF activity, Ch^WT^ seedlings showed an etiolated phenotype similar to that of At^WT^ when grown in the dark, in contrast to *A. thaliana pifq* and 35S:GFP-ΔNt-HFR1 seedlings (**Figure 6g**), suggesting the PIF activity is high enough in *C. hirsuta* to induce the normal skotomorphogenic development.

## DISCUSSION

It is currently unknown whether the switch between shade avoidance and tolerance strategies is an easily adjustable trait in plants. The existence of closely related species with divergent strategies to acclimate to shade provides a good opportunity to study the genetic and molecular basis for adapting to this environmental cue. To this goal, we performed comparative analyses of the hypocotyl response to shade in young seedlings of two related *Brassicaceae*: *A. thaliana* and *C. hirsuta. A. thaliana*, a model broadly used to study the SAS hypocotyl response, is well characterized on a physiological, genetic and molecular level. *C. hirsuta* was previously described as a shade tolerant species whose hypocotyls are unresponsive to simulated shade ^25,26^. Recent work showed that phyA is a major contributor to the suppression of hypocotyl elongation of *C. hirsuta* seedlings in response to shade, mainly due to the stronger phyA activity in this species compared to the shade-avoider *A. thaliana* ^26^. Importantly, an enhanced phyA activity was not enough to explain the lack of shade-induced hypocotyl elongation in *C. hirsuta*, pointing to additional components that contribute to this response. Our aim to fill this gap led us to uncover a role for HFR1 in this response.

In *C. hirsuta*, removal of HFR1 function resulted in a strong *slender in shade* (*sis*) phenotype but milder than that of *sis1* plants, deficient in the phyA photoreceptor ^26^, providing genetic evidence for the role of *HFR1* in restraining the *C. hirsuta* hypocotyl elongation in shade (**Figure 1a, b**). This indicates that, like phyA, HFR1 contributes to implement the shade tolerant habit in *C. hirsuta* seedlings. Because of the *sis* phenotype of *chfr1* and RNAi-HFR1 seedlings (**Figure 1**) we hypothesized that HFR1 activity is higher in *C. hirsuta* than in *A. thaliana*. Consistently, transcript levels of *HFR1* were significantly higher in Ch^WT^ than At^WT^ seedlings in both W and W+FR (**Figure 2**). Higher HFR1 levels in *C. hirsuta* may not be relevant in W because of the expected lower abundance and activity of PIFs, but a higher pool of ChHFR1 ready to suppress early ChPIF action in shade could provide a fast and sustained repression of the elongation response. Indeed, the shade-induced expression of *PIL1, YUC8* and *XTR7*, known to be direct PIF target genes in *A. thaliana*, was strongly and rapidly enhanced in *chfr1* and RNAi-HFR1 seedlings (**Figure 1c, d**). More importantly, rapid shade-induced expression was globally attenuated in Ch^WT^ compared to At^WT^ seedlings ^26^.

In addition to changes in gene expression, a higher HFR1 activity in *C. hirsuta* could also result from post-translational regulation affecting protein stability. Our immunoblot analyses indicated that HFR1 proteins rapidly accumulate in response to simulated shade (W+FR), likely as a consequence of the strong shade-induced responsiveness of the promoter (**Figure 4a**). These results support that regulation of HFR1 protein abundance in low R:FR occurs mainly at the transcriptional level, as suggested ^2^. More importantly, ChHFR1 accumulates significantly more when under the control of a constitutive promoter either under W or W+FR and it is degraded at a slower pace than AtHFR1 in shade (**Figure 4b, c**) suggesting that intrinsic differences in post-translational stability between these proteins play a role in their contrasting accumulation.

AtHFR1 protein abundance is modified post-translationally by phosphorylation ^41^ and ubiquitination in a light-dependent manner ^31,42^. Canopy shade promotes nuclear accumulation of COP1 ^43,44^ allowing it to directly interact with and polyubiquitinate AtHFR1, leading to its degradation by the 26S proteasome ^31,42,45^. AtHFR1 contains two putative COP1 binding sites (amino acids 48-83) on its N-terminal domain (Nt, amino acids 1-131), although only one site has been shown to bind COP1 ^46^. Deletion of AtHFR1 Nt leads to its stabilization in the dark and light ^30^, and results in a stronger biological activity ^19,31,42^, highlighting the importance of the COP1-interacting domain for light regulation of AtHFR1. AtHFR1 and ChHFR1 primary structures are similar, including the putative COP1-interacting domain ^42^, except for the addition of 30 amino acids at the N-terminal part of ChHFR1 and a 9-amino acid insertion in the C-terminal part of AtHFR1 (**Supplemental Figure 4a**). Therefore, protein sequence and/or other structural differences (e.g., COP1 binding sites) between AtHFR1 and ChHFR1 (**Supplemental Figure 4**) could influence their differential stability and, at least in part, may account for the difference in response to shade between *C. hirsuta* and *A. thaliana*.

AtHFR1 was previously shown to interact with all the AtPIFQ members and to form non-DNA-binding heterodimers ^20-22^. Our genetic and Y2H experiments extended the list of AtHFR1 interactors to AtPIF7, the major SAS-promoting PIF (**Figure 5**). If ChHFR1 maintains similar PIF-binding abilities, the reduced expression of *ChPIF7* (**Figure 2**) might further contribute to imbalance the PIF-HFR1 module in favor of the negative HFR1 activity in *C. hirsuta* compared to *A. thaliana*. Because of the higher stability of ChHFR1 over AtHFR1 in shade (**Figure 4**), an even stronger repression of global PIF activity in *C. hirsuta* would contribute to the unresponsiveness of hypocotyls to shade. The attenuation of the warm temperature-induced hypocotyl elongation in *C. hirsuta*, which is a PIF-regulated process in *A. thaliana* ^36,37,47,48^ and HFR1-dependent in both species (**Figures 6a-c**), further agrees with our proposal of an enhanced activity of HFR1 in *C. hirsuta* compared to *A. thaliana*. On the other hand, the delayed DIS observed in *C. hirsuta*, shown to be PIF-regulated in *A. thaliana* ^38,40^ but unaffected by HFR1 in the two species analyzed (**Figures 6d, e**), suggests that PIF activity is globally lower *per se* in *C. hirsuta* than in *A. thaliana*. Together, our results indicate that PIF-HFR1 module is imbalanced in *C. hirsuta* by the combination of (1) an attenuated *PIF7* expression compared to *A. thaliana*, and (2) the increased levels of ChHFR1 in light and shade conditions, resulting in the repression of PIF-regulated processes in *C. hirsuta* (**Figure 7**). Importantly, although attenuated, PIF activity in *C. hirsuta* is enough to provide a functional and effective etiolation response (**Figure 6g**) for seedlings survival during germination in the dark.

**Figure 7.**
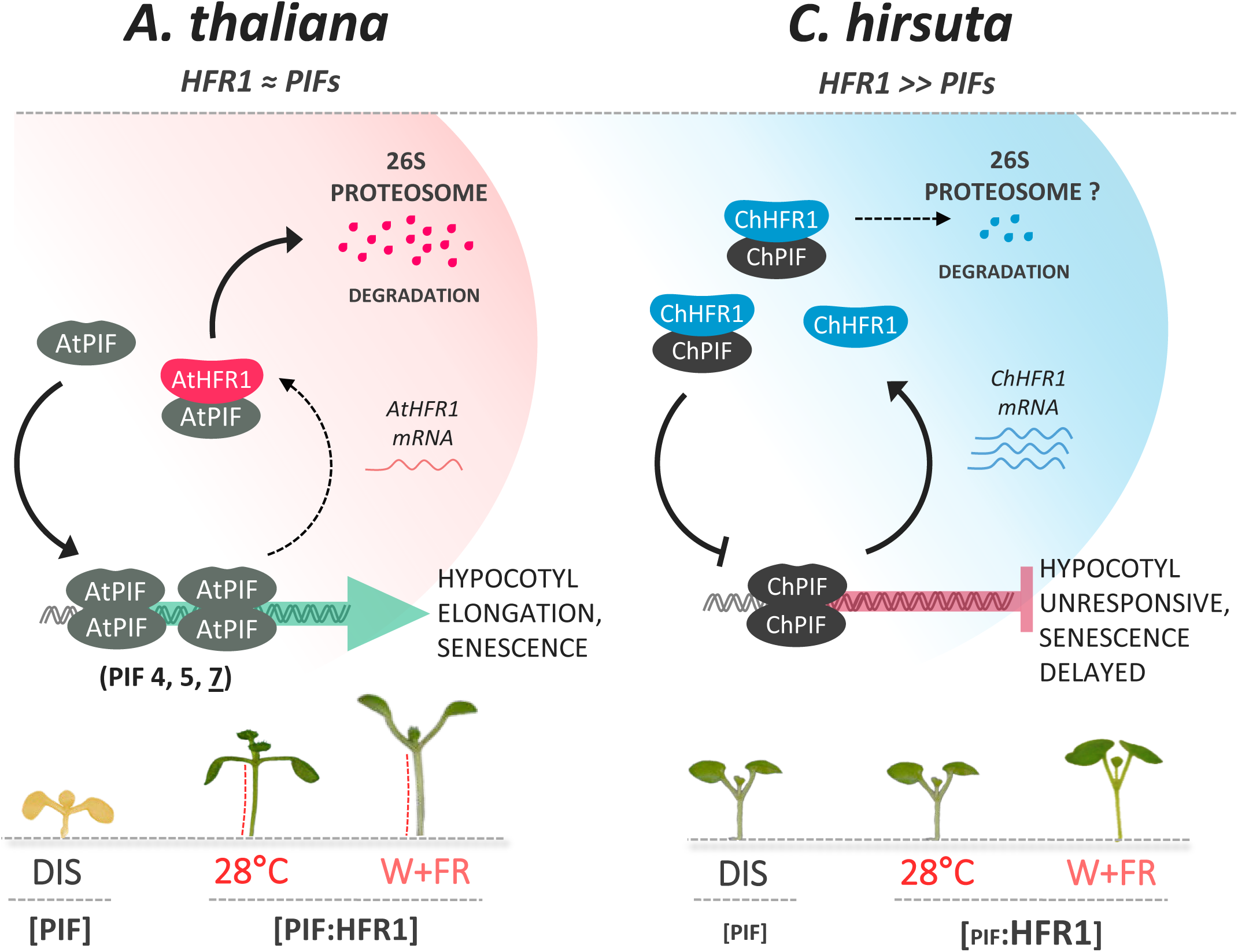
Model summarizing how PIF-HFR1 transcriptional module is differently balanced in *A. thaliana* and *C. hirsuta*. Shade (low R:FR) displaces phytochrome photoequilibrium towards the inactive form, allowing PIFs to promote the expression of shade avoidance related genes, such as *HFR1*. PIF transcript or/and protein levels are induced in response to warm temperatures, resulting in enhanced expression of growth-promoting genes. HFR1 abundance is also increased by warm temperature. HFR1 modulates these responses by heterodimerizing with PIFs and inhibiting their DNA-binding ability. As a result, HFR1 attenuates hypocotyl elongation of *A. thaliana* seedlings in response to shade or warm temperature. In *C. hirsuta*, higher HFR1 activity inhibits more effectively PIF action than in *A. thaliana*. In addition, PIF abundance is attenuated in *C. hirsuta*. Both changes alter the PIF-HFR1 balance in *C. hirsuta*, resulting in lower PIF transcriptional activity. As a consequence, shade- and warm temperature-induced hypocotyl elongation are repressed and DIS is delayed in this species.

Activity of HFR1 and phyA ^26^ appears to be increased in *C. hirsuta* to maintain unresponsiveness of hypocotyls to shade. An aspect shared by both negative regulators is that their expression and/or stability are strongly affected by light conditions. Expression of both *PHYA* and *HFR1* is induced by simulated shade in de-etiolated seedlings. By contrast, whereas the stability of the photolabile phyA is reduced by light but enhanced by shade, that of AtHFR1 is promoted by light and decreased by shade ^27,30,41,43,49-52^. Although expression of both *PHYA* and *HFR1* is higher in *C. hirsuta* than in *A. thaliana*, different mechanisms might contribute to their increased activity in *C. hirsuta*. Indeed, enhanced ChphyA repression was achieved by its stronger specific intrinsic activity ^26^. By contrast, enhanced ChHFR1 repression was accomplished through its higher gene expression and protein stability coupled with an attenuated PIF7 activity. Altogether this could provide a more repressive state of the *C. hirsuta* PIF-HFR1 module. Because of the temporal differences downregulating many of the shade marker genes between phyA (observed after 4-8 hours of shade exposure) ^26^ and HFR1 (rapidly detected after just 1 h of shade exposure) (**Figure 1c, d**), it seems likely that ChHFR1 and ChphyA suppressor mechanisms of shade response in *C. hirsuta* act independently, as it was reported for *A. thaliana* ^27,53^. Therefore, the concerted activity of these two independent suppressor mechanisms seems to coordinately prevent the shade-induced hypocotyl elongation in *C. hirsuta*. Whether other shade tolerant species employ the same adaptive principles is something we aim to explore in the future.

## METHODS

### Plant material and growth conditions

*Arabidopsis thaliana hfr1-5, pif7-1, pif7-2* and *pifq* mutants, 35S:PIF7-CFP and 35S:GFP-ΔNt-HFR1 lines (in the Col-0 background, At^WT^) and *Cardamine hirsuta* (Oxford ecotype, Ox, Ch^WT^) plants have been described before ^19,25,31,33^. Plants were grown in the greenhouse under long-day photoperiods (16 h light and 8 h dark) to produce seeds, as described ^14,50,54^. For transient expression assays, *Nicotiana benthamiana* plants were grown in the greenhouse under long-day photoperiods (16 h light and 8 h dark).

For hypocotyl assays, seeds were surface-sterilized and sown on solid growth medium without sucrose (0.5xGM–). For gene expression analyses, immunoblot experiments and pigment quantification, seeds were sown on a sterilized nylon membrane placed on top of the solid 0.5xGM– medium. After stratification (dark at 4°C) of 3-6 days, plates with seeds were incubated in plant chambers at 22°C under continuous white light (W) for at least 2 h to break dormancy and synchronize germination ^55,56^.

W was emitted from cool fluorescent tubes that provided from 20 to 100 µmol m^-2^ s^-1^ of photosynthetically active radiation (PAR) with a red (R) to far-red light (FR) ratio (R:FR) from 1.3-3.3. The different simulated shade treatments were produced by supplementing W with increasing amounts of FR (W+FR). FR was emitted from GreenPower LED module HF far-red (Philips), providing R:FR of 0.02-0.09. Light fluence rates were measured with a Spectrosense2 meter (Skye Instruments Ltd) ^50^.

Temperature induced hypocotyl elongation assays were done by placing the plates with seeds under the indicated light conditions in growth chambers at 22°C or 28°C.

### Measurement of hypocotyl length

Hypocotyl length was measured as described ^55,56^. Experiments were repeated at least three times with more than 10 seedlings per genotype and/or treatment, and average values are shown.

### Generation of transgenic lines, mutants and crosses

*A. thaliana hfr1-5* plants were transformed to express *AtHFR1* and *ChHFR1* under the promoters of 35S or *AtHFR1* (pAt). The obtained lines were named as *hfr1>35S:ChHFR1, hfr1>pAt:AtHFR1* and *hfr1>pAt:ChHFR1*. Transgenic RNAi-HFR1 and mutant *chfr1-1* and *chfr1-2* lines are in Ch^WT^ background. Details of the constructs used for the generation of these lines ^57^ are provided as Supplementary information.

### Gene expression analyses

Real-time qPCR analyses were performed using biological triplicates, as indicated ^14^. Total RNA was extracted from seedlings, treated as indicated, using commercial kits (Maxwell® SimplyRNA and Maxwell® RSC Plant RNA Kits; www.promega.com). 2 µg of RNA was reverse-transcribed with Transcriptor First Strand cDNA synthesis Kit (Roche, www.roche.com). The *A. thaliana UBIQUITIN 10* (*UBQ10*) was used for normalization in *A. thaliana hfr1-5* lines expressing *AtHFR1* or *ChHFR1*. The *ELONGATION FACTOR 1α* (*EF1α*), *YELLOW-LEAF-SPECIFIC GENE 8* (*YLS8*) and *SPC25* (AT2G39960) were used for normalizing and comparing the levels of *HFR1* and *PIF7* between *A. thaliana* and *C. hirsuta*. All primers sequences for qPCR analyses are provided as Supplementary information (**Supplemental Table 1**).

### Protein extraction and immunoblotting analyses

To detect and quantify transgenic AtHFR1 and ChHFR1, proteins were extracted from ∼50 mg of 7-day old seedlings (grown as indicated) or from 50-75 mg of agroinfiltrated *N. benthamiana* leaves. Plant material was frozen in liquid nitrogen, ground to powder and total proteins were extracted using an SDS-containing extraction buffer (1.5 µL per mg of fresh weight), as described ^14^. Protein concentration was estimated using Pierce™ BCA Protein Assay Kit (Thermo Scientific, www.thermofisher.com). Proteins (45 - 50 µg per lane) were resolved on a 10% SDS-PAGE gel, transferred to a PVDF membrane and immunoblotted with rat monoclonal anti-HA (High Affinity, clone 3F10, Roche, www.roche.com; 1:2000 dilution) and hybridized with peroxidase conjugated goat anti-rat (Polyclonal, A9037, Sigma, www.sigmaaldrich.com; 1:5000 dilution) and after membrane stripping, with rabbit polyclonal anti-actin (Agrisera, www.agrisera.com; 1:5000 dilution) then hybridized with peroxidase conjugated donkey anti-rabbit (Amersham, www.gelifesciences.com; 1:10000 dilution). Development of blots was carried out in ChemiDoc™ Touch Imaging System (Bio-Rad, www.bio-rad.com) using ECL Prime Western Blotting Detection Reagent (GE Healthcare, RPN2236). Relative protein levels of three to four biological replicates were quantified using Image Lab™ Software (Bio-Rad, www.bio-rad.com).

### Yeast 2 Hybrid (Y2H) assays

For Y2H assays we employed a cell mating system, as described ^14^. The leucine (Leu) auxotroph YM4271a yeast strain was transformed with the AD-derived constructs and the tryptophan (Trp) auxotroph pJ694α strain with the BD-derived constructs. Colonies were selected on synthetic defined medium (SD) lacking Leu (SD-L) or Trp (SD-W), grown in liquid medium and set to mate by mixing equal volumes of transformed cells. Dilutions of the mated cells were selected on SD-LW and protein interactions were tested on SD-LW medium lacking histidine (SD-HLW). Details of the yeast constructs used are provided as Supplementary information.

## ACKNOWLEDGEMENTS

We are grateful to Peter Quail (PGEC, Albany, CA, USA) for providing 35S:PIF7-CFP seeds; and to Manuel Rodriguez-Concepción (CRAG) for comments on the manuscript. SP received predoctoral fellowships from the *Agència d’Ajuts Universitaris i de Recerca* (AGAUR - Generalitat de Catalunya, FI program). CT received a Marie Curie IEF postdoctoral contract funded by the European Commission and a CRAG short-term fellowship. Our research is supported by grants from BBSRC (BB/H006974/1) and Max Planck Society (core grant) to MT, and from MINECO-FEDER (BIO2017-85316-R, and BIO2017-84041-P) and AGAUR (2017-SGR1211, 2017-SGR710 and Xarba) to JFM-G and MRC. We also acknowledge the support of the MINECO for the “Centro de Excelencia Severo Ochoa 2016-2019” award SEV-2015-0533 and by the CERCA Programme / Generalitat de Catalunya. The IJPB benefits from the support of the LabEx Saclay Plant Sciences-SPS (ANR-10-LABX-0040-SPS).

