## Supplemental information for "Adaptation to plant shade relies on rebalancing the transcriptional activity of the PIF-HFR1 regulatory module"

### SUPPLEMENTAL METHODS

#### Generation of RNAi-HFR1 plants of *C. hirsuta*

To generate an RNAi construct for silencing of the endogenous *ChHFR1*, a fragment of 222 bp was PCR amplified using primers CTO35 + CTO36 (Supplemental Table 2) and cDNA of 7-day old *C. hirsuta* seedlings grown 1 h under W+FR. This partial fragment of *ChHFR1* (ptChHFR1) was cloned into pCRII-TOPO (Invitrogen, www.thermofisher.com) to generate pCT17, which was confirmed by sequencing. An *Eco*RI fragment of pCT17 was subcloned into pENTR3C vector (Invitrogen, www.thermofisher.com), to create the Gateway entry clone pCT19 (to have ptChHFR1 flanked with attL1 and attL2, attL1<ptChHFR1<attL2). Recombination of pCT19 with the destination vector pB7GWIWG2(I), which contains attR1 and attR2 sites, using Gateway LR Clonase II (Invitrogen), gave pCT33 (35S:attB1<RNAi-ChHFR1<attB2). This plasmid is a binary vector conferring resistance to the herbicide phosphinothricin (PPT) in plants and the antibiotic spectinomycin in bacteria. *Agrobacterium tumefaciens* strain $C_{58}C_1$ (pGV2260) was transformed with pCT33 by electroporation and colonies were selected on solid YEB medium with rifampicin (100 µg/mL),

kanamycin (25 µg/mL) and spectinomycin (100 µg/mL). Wild-type *C. hirsuta* (Ox, Ch$^{WT}$) plants were transformed by floral dipping and transgenic seedlings were selected on 0.5xGM- medium [1] containing 50 µg/mL PPT. Transgene in seedlings of T1 generation was verified by PCR genotyping using specific primers. Plants homozygous for the transgene were finally used for experiments.

### Isolation of *HFR1* mutants of *C. hirsuta*

To obtain loss-of-function mutants of *ChHFR1* in *C. hirsuta* (named as *chfr1*) we employed the CRISPR-Cas9 gene editing system [2]. The guide RNA targeting ChHFR1 (gRNA$_{ChHFR1}$, 5'-GTT-GAA-GAC-TGC-AGA-TTT-GT-3') was synthesized to be under the control of the *A. thaliana U6* promoter (pU6) sequence and flanked by the Gateway attB1 and attB2 recombination sites (IDT, https://eu.idtdna.com/pages) (attB1<pU6:gRNA$_{ChHFR1}$<attB2). This sequence was recombined with the vector pDONR207 using Gateway BP Clonase II (Invitrogen) to generate the entry vector pSP101 (attL1<pU6:gRNA$_{ChHFR1}$<attL2). In a recombination reaction of pSP101 with pDE-Cas9 [3] using Gateway LR Clonase II (Invitrogen), a binary vector pSP102 was created (attB1<pU6:gRNA$_{ChHFR1}$<attB2, Cas9). This vector, that contains the information to target ChHFR1, confers resistance to PPT in plants and spectinomycin in bacteria. *A. tumefaciens* strain C$_{58}$C$_{1}$ (pGV2260) was transformed with pSP102 by electroporation and colonies were selected on solid YEB medium with antibiotics, as indicated before for pCT33. Wild-type *C. hirsuta* (Ox, Ch$^{WT}$) plants were transformed by floral dipping and resistant transgenic seedlings were selected on 0.5xGM- medium containing PPT (30 µg/mL). These T1 seedlings were PCR genotyped using primers MJO27 and

MJO28 (Supplemental Table 2) to detect the presence of the transgene. In the following T2 generation, a total of six seedlings with a *sis* phenotype from 1 independent transgenic line were selected and grown to maturity. An *HFR1* fragment of 664 bp around the $gRNA_{ChHFR1}$ target sequence was amplified by PCR from gDNA of each plant using primers CTO29 + CTO36 (Supplemental Table 2). Sequencing of these fragments indicated the presence of mutations in *ChHFR1* gene. Descendants of these plants (T3 generation) were reselected in shade and sequenced to confirm the unambiguous presence of the mutated *chfr1* alleles. In the T4 generation, seedlings sensitive to PPT (indicating the loss of T-DNA insertion) were selected, which resulted in the isolation of the *chfr1-1* and *chfr1-2* mutant allele lines (Figure S1). These mutants were genotyped by PCR using primers SPO104 + SPO107 (for *chfr1-1*) and SPO106 + SPO107 (for *chfr1-2*) (Supplemental Table 2).

**Generation of *A. thaliana hfr1-5* transgenic lines expressing *AtHFR1* or *ChHFR1* under the control of different promoters**

We amplified a 2 kbp fragment of *AtHFR1* promoter starting immediately before the ATG of *AtHFR1* gene using gDNA of *A. thaliana* wild-type Col-0 ($At^{WT}$) as a template and primers SPO26 + SPO27 (Table S2). This fragment was subcloned into pCRII-TOPO (Invitrogen) to generate pSP51. From the different clones analyzed, the best one was pSP51.10, with three 1 bp-deletions in the amplified region, none affecting the G-boxes, known to be necessary for PIF binding.

*AtHFR1* coding sequence was amplified from pJB30 [4] using primers RO25 + SPO30 (Supplemental Table 2), which removed the stop codon and introduced a *Xho*I site at the N-terminal site. After subcloning this fragment into pCRII-TOPO, which gave pSP54 (*AtHFR1*), the insert was sequenced to confirm its identity. The 3xHA fragment was amplified from plasmid pEN-R2-3xHA-L3 [5] and primers SPO31 (which added a *Sal*I site) + SPO32 (which added a *Xho*I site, Supplemental Table 2). This fragment was subcloned into pCRII-TOPO to generate pSP55 (*3xHA*), whose insert was sequenced to confirm its identity. A *BamH*I-*Xho*I fragment of pSP54 was subcloned into pSP55 digested with *BamH*I and *Sal*I to generate pSP57 (*AtHFR1-3xHA*). A *BamH*I-*Xho*I fragment of pSP57 was subcloned into the same sites of pENTR3C vector (Invitrogen) which gave pSP59. This plasmid contained *AtHFR1-3xHA*, with an extra *Xba*I site in the C-terminus end, flanked with attL1 and attL2 sites (attL1<*AtHFR1-3xHA*$^{XbaI}$<attL2). *Xba*I restriction site in pSP59 was removed by filling the site with Klenow enzyme after digestion, and religation to generate pSP84 (attL1<*AtHFR1-3xHA*<attL2). Recombination of pSP84 with the binary vector pIR101 (attR1<*ccdB*<attR2) [6] (using Gateway LR Clonase II (Invitrogen) resulted in pSP88 (attB1<*AtHFR1-3xHA*<attB2). An *Xba*I fragment of pSP51 was subcloned into the same site of pSP88 which gave pSP90 (*pAtHFR1*:attB1<*AtHFR1-3xHA*<attB2). This binary vector confers resistance to spectinomycin in bacteria and PPT in plants.

*ChHFR1* CDS was amplified using cDNA from wild-type *C. hirsuta* (Ox, Ch$^{WT}$) seedlings and primers SPO28 + SPO29 (Supplemental Table 2), which removed the stop codon and introduced a *Xho*I site. This PCR product was subcloned into pCRII-TOPO to generate pSP53 (*ChHFR1*). Selected colonies were sequenced to confirm their identity. A *BamH*I-*Xho*I fragment of pSP53 was subcloned into pSP55 digested with *BamH*I-*Sal*I to generate pSP56 (*ChHFR1-3xHA*). A *BamH*I-*Xho*I fragment of pSP56 was subcloned into the same site of pENTR3C vector (Invitrogen), which gave pSP58. This plasmid contained *ChHFR1-3xHA*, with an *Xba*I site in the C-terminus end, flanked with attL1 and attL2 sites (attL1<*ChHFR1-3xHA*$^{XbaI}$<attL2). *Xba*I restriction site in pSP58 was removed by filling the site with Klenow enzyme after digestion, and religation to generate pSP83 (attL1<*ChHFR1-3xHA*<attL2). Recombination of pSP83 with the binary vector pIR101 using Gateway LR Clonase II (Invitrogen) resulted in pSP87 (attB1<*ChHFR1-3xHA*<attB2). An *Xba*I fragment of pSP51 was subcloned into the same site of pSP87 which gave pSP89 (*pAtHFR1*:attB1<*ChHFR1-3xHA*<attB2). This binary vector confers resistance to spectinomycin in bacteria and PPT in plants.

To overexpress *ChHFR1*, a *BamH*I-*Xho*I fragment of pSP58 was subcloned into the *BamH*I-*Sal*I digested pCAMBIA1300 based pCS14 [7] to generate pSP81 (*35S:ChHFR1*). This binary vector confers resistance to kanamycin in bacteria and hygromycin in plants.

*A. thaliana hfr1-5* plants were transformed with pSP81, pSP89 and pSP90, as previously described. Transgenic seedlings were selected on 0.5xGM- medium with PPT (15 µg/mL) or hygromycin (30 µg/mL), verified by PCR genotyping using specific primers. Homozygous transgenic plants with 1 T-DNA insertion were finally used for experiments.

**Generation of constructs for transient expression in *N. benthamiana* leaves**

To overexpress *ChHFR1* and *AtHFR1* in *N. benthamiana*, a Gateway vector was created using pCAMBIA1302 (*35S:mGFP5*) as a backbone. An *Nsi*I-*Hind*III fragment of pEarlyGate 100 (*35S*:attR1<*ccdB*<attR2) [8] was subcloned into pCAMBIA1302 digested with *Pst*I-*Hind*III, which gave pSP135 (*35S*:attR1<*ccdB*<attR2, *35S:mGFP5*). Recombination of pSP58 and pSP59 (both linearized with *Nhe*I) with the binary vector pSP135 using Gateway LR Clonase II (Invitrogen) gave pSP141 (*35S*:attB1<*ChHFR1-3xHA*<attB2, *35S:mGFP5*) and pSP142 (*35S*:attB1<At*HFR1-3xHA*<attB2, *35S:mGFP5*), respectively. These two binary vectors also overexpress mGFP5 and confer resistance to kanamycin in bacteria.

*N. benthamiana* plants were agroinfiltrated with pSP141 or pSP142, and the strain expressing the HcPro protein [9]. Samples were taken 3 days after agroinfiltration.

**Generation of constructs for the Yeast 2 Hybrid (Y2H) assays**

*AtPIF7* CDS was amplified using cDNA of *A. thaliana* wild-type Col-0 (At$^{WT}$) seedlings and primers JO414 + JO415 (Supplementary Table 2), which removed the STOP codon and introduced a *Xho*I site. This PCR product was subcloned into pCRII-TOPO to generate pRA1 (*AtPIF7*). The insert was sequenced to confirm its identity. A *Xho*I fragment of pRA1 was subcloned into pSP55 digested with *Sal*I to generate pRA2 (*AtPIF7-3xHA*). An *Eco*RI fragment of pRA2 was subcloned into the same site of pENTR3C entry vector (Invitrogen) which gave pRA3 (attL1<*AtPIF7-3xHA*<attL2). This PIF7-3xHA had a stop codon immediately before the ATG, which prevented from cloning it in frame with the yeast derived proteins.

Therefore, the *PIF7-3xHA* gene was PCR amplified using pRA3 as a DNA template and primers BAO4 + BAO5 (Supplemental Table 2) to add attB1 and attB2 sequences (attB1<*AtPIF7-3xHA*<attB2). This fragment was recombined with pDONR207 using Gateway BP Clonase II (Invitrogen) to obtain pBA7 (attL1<*AtPIF7-3xHA*<attL2). The insert was sequenced to confirm its identity. In a recombination reaction of pBA7 and pGBKT7-GW [10] which contained the Gal4 DNA-binding domain (BD, attR1<ccdB<attL2; it confers Trp auxtrophy), and pBA7 and pGADT7-GW [10] which contained the Gal4 activation domain (AD, attR1<ccdB<attL2; it confers Leu auxtrophy), using Gateway LR Clonase II (Invitrogen), pBA10 (BD-attB1<*AtPIF7-3xHA*<attB2) and pBA11 (AD-attB1<*AtPIF7-3xHA*<attB2) were obtained. These plasmids allowed expressing the fusion BD-PIF7-3xHA or AD-PIF7-3xHA proteins under the *ADH1* promoter in yeast, respectively.

**GUS lines**

Transgenic lines expressing GUS were based on a modified pIR101 plasmid [6] which contains the reporter *GUS* gene in a promoterless context (attB1<*GUS*<attB2). *Xba*I fragment of pSP51 was subcloned into the same site of modified pIR101 to give pSP86 (pAtHFR1:attB1<*GUS*<attB2). This binary vector confers resistance to spectinomycin in bacteria and PPT in plants. *A. thaliana* wild-type Col-0 ($At^{WT}$) plants were transformed with this construct as described previously.

**GUS staining**

Histochemical GUS assays were done as described [1], incubating seedlings at 37ºC without ferricyanide/ferrocyanide.

### SUPPLEMENTAL FIGURE LEGENDS

**Supplemental Figure 1. Characterization of RNAi-HFR1 and *chfr1* mutants in *C. hirsuta*.** Relative expression levels of *ChHFR1* gene, normalized to *EF1α* in $Ch^{WT}$, **(a)** two RNAi-HFR1 lines (#01 and #21) and **(b)** the two *chfr1* mutants of *C. hirsuta.* Seedlings were grown for 7 days in W. Expression values are the mean ± SE of three independent biological replicates relative to $Ch^{WT}$. **(c)** The two identified *chfr1-1* and *chfr1-2* mutants have one nucleotide insertion at position 420 of the *ChHFR1* ORF, which leads to a frame shift and a premature stop codon.

**Supplemental Figure 2. Alignments of *HFR1*, *PIF4*, *PIF5* and *PIF7* partial DNA sequences in *A. thaliana* and *C. hirsuta* (a).** Location of shared primers and amplicons used for comparison of expression levels by RT-qPCR between species. **(b)** Transcript abundance of *PIF4* and *PIF5*, normalized to *YLS8*, *SPC25* and *EF1α* in $Ch^{WT}$ and $At^{WT}$ grown as in Figure 2. Expression values are the means ± SE of three independent biological replicates relative to the data of $At^{WT}$ grown in continuous W at day 3. Asterisks mark significant differences (2-way ANOVA: * p-value <0.05, ** p-value <0.01, *** p-value <0.001) between $Ch^{WT}$ and $At^{WT}$ when grown under W (black asterisks) or W+FR (red asterisks).

**Supplemental Figure 3. The 2 Kbp *AtHFR1* promoter is shade responsive in *A. thaliana*. (a)** Cartoon of *HFR1* promoters from *A. thaliana* (*pAtHFR1*) and *C. hirsuta* (*pChHFR1*). These promoters cover 2000 bp from the beginning of the translation start of the two *HFR1* genes. The positions of G-boxes (CACGTG) are indicated with arrows. **(b)** GUS staining of representative *A. thaliana* seedlings expressing *GUS* under the *pAtHFR1* (line #03). Seven-day-old W-grown seedlings were treated with W+FR for the indicated amount of time.

**Supplemental Figure 4. ChHFR1 protein accumulates in high W. (a)** Alignment of AtHFR1 and ChHFR1 protein sequences. Putative COP1 interacting motifs, defined in AtHFR1, are indicated in blue. **(b)** Cartoon representing the light treatments given to seedlings to estimate relative HFR1-3xHA levels. Seedlings grown for 7 d in low W (~20 μmol $m^{-2}$ $s^{-1}$, R:FR≈6.4) were first moved to high W (~100 μmol $m^{-2}$ $s^{-1}$, R:FR≈3.9) for 3 h and then either transferred to high W (control) or high W+FR (R:FR≈0.06) for 3 h. Seedling samples were collected at the time points indicated with asterisks. **(c)** Relative HFR1-3xHA protein levels of *hfr1>35S:ChHFR1* seedlings (line #16) grown as indicated in **b**, with a representative immunoblot in a lower panel. Relative protein levels are the mean ± SE of three independent biological replicates relative to the data point of 0 h in high W (0 h W). Asterisks mark significant differences in protein levels (Student *t*-test: ** p-value <0.01; * p-value <0.05) relative to the 0 h W value.

**Supplemental Figure 5. Relative expression levels of *AtHFR1* and *AtPIF7* genes in transgenic lines overexpressing *GFP-ΔNt-HFR1* and/or *PIF7-CFP*.** Relative expression, normalized to *UBQ10*, was estimated in seedlings grown for 7 days in W. Expression values are the mean ± SE of three independent biological replicates relative to $At^{WT}$. Asterisks mark significant differences in expression (Student *t*-test: ** p-value <0.01; * p-value <0.05) relative to 35S:GFP-ΔNt-HFR1-GFP or 35S:PIF7-CFP values.

### SUPPLEMENTAL TABLES:

**Supplemental Table 1. Primers used for gene expression analyses.** Primers BO40 and BO41 for amplifying *UBQ10* (Sorin *et al.*, 2009), SPO102 and SPO103 (*AtEF1α* and *ChEF1α*), SPO113 and SPO114 (*AtSPC25* and *ChSPC25*), and SPO115 and SPO116 (*AtYLS8* and *ChYLS8*) have been described before [6].

| Gene | Primer name | Sequence (5' – 3') |
|---|---|---|
| *ChEF1α* | CTO9 (F) | GGCCGATTGTGCTGTCCTTA |
| | CTO10 (R) | TCACGGGTCTGACCATCCTTA |
| *ChHFR1* | CTO13 (F) | CGGCGTCGTGTCCAGATC |
| | CTO14 (R) | TGAACCTTTTCGCGTCAGTG |
| *ChPIL1* | CTO17 (F) | GAAGACCCCAAAACAACGGTT |
| | CTO18 (R) | CCCTCATCGTACTCGGTCTCA |
| *ChYUC8* | CTO51 (F) | TTACGCCGGGAAAAAAGTTCT |
| | CTO52 (R) | GCGAAATGGTTGGCTAGGTC |
| *ChXTR7* | CTO69 (F) | TGGTGTTCCTTTCCCAAAAAA |
| | CTO70 (R) | CCACCTCTCGTAGCCCAATC |
| *AtHFR1, ChHFR1* | SPO88 (F) | CCAGCTTCTTCTCCTCA |
| | SPO89 (R) | CATCGCATGGGAAGAAAAATC |
| *AtPIF4, ChPIF4* | SPO108 (F) | CCAATACCCTCCAGATGAAGAC |
| | SPO109 (R) | TCTCTGAGGTTGGTCTCTGG |
| *AtPIF5, ChPIF5* | SPO110 (F) | CATTAATCAGATGGCTATGCA |
| | SPO111 (R) | AACTGTACCGGGTTTTGACA |
| *AtPIF7, ChPIF7* | SPO112 (F) | TCCGCTCTGGATCGGAAACTC |
| | SPO64 (R) | TGCTCGTCCCCGTCGTCCAT |
| | SPO142 (R) | TCTCATCCTCTGGTTTATCC |

**Supplemental Table 2. Primers used for cloning or/and genotyping.** Primer RO25 [11] has been described before.

| **Gene** | **Primer name** | **Sequence (5' – 3')** |
|---|---|---|
| *ChHFR1* WT | SPO104 (F) | CTGTTGAAGACTGCAGATTTG |
| | SPO107 (R) | CCTAAGGCAAGATTCTTTGAA |
| *chfr1-1* | SPO105 (F) | CTGTTGAAGACTGCAGATTA |
| *chfr1-2* | SPO106 (F) | CTGTTGAAGACTGCAGATTTT |
| attB1 | MJO27 (F) | GGGGACAAGTTTGTACAAAAAAGCAGGCT |
| attB2 | MJO28 (R) | GGGGACCACTTTGTACAAGAAAGCTGGGT |
| *pAtHFR1* | SPO26 (F) | GCTCTAGAGTAAAGATAACGTTCT |
| | SPO27 (R) | GCTCTAGAGTTAGTTAAAGAGATA |
| *ChHFR1* | SPO28 | CCATGGGTTTTCCATTTTCTCG |
| | SPO29 (R) | GGCTCGAGGAGTCTTCCCATCGCA |
| *ChHFR1* | CTO29 (F) | ATGATCATCATCAAATTGTTC |
| *AtHFR1* | SPO30 (R) | GGCTCGAGTAGTCTTCTCATCGCA |
| 3xHA | SPO31 (F) | CCGTCGACGGTGGAGGCGGTTCAG |
| | SPO32 (R) | GGCTCGAGTCAAGCGTAATCTGGA |
| RNAi-ChHFR1 | CTO35 (F) | CAAACACATAATGATCATCATC |
| | CTO36 (R) | ATCACTCCAGATCTGGACACGA |
| *AtPIF7* | JO414 (F) | TAACACATGTCGAATTATGGAG |
| | JO415 (R) | GGCTCGAGATCTCTTTTCTCATGATTC |
| *AtPIF7* + attB1 | BAO4 (F) | GGGGACAAGTTTGTACAAAAAAGCAGGCTAC ATGTCGAATTATGGAGTTAAAG |
| *AtPIF7* + attB2 | BAO5 (R) | GGGGACCACTTTGTACAAGAAAGCTGGGTGT CAAGCGTAATCTGGAACGTC |

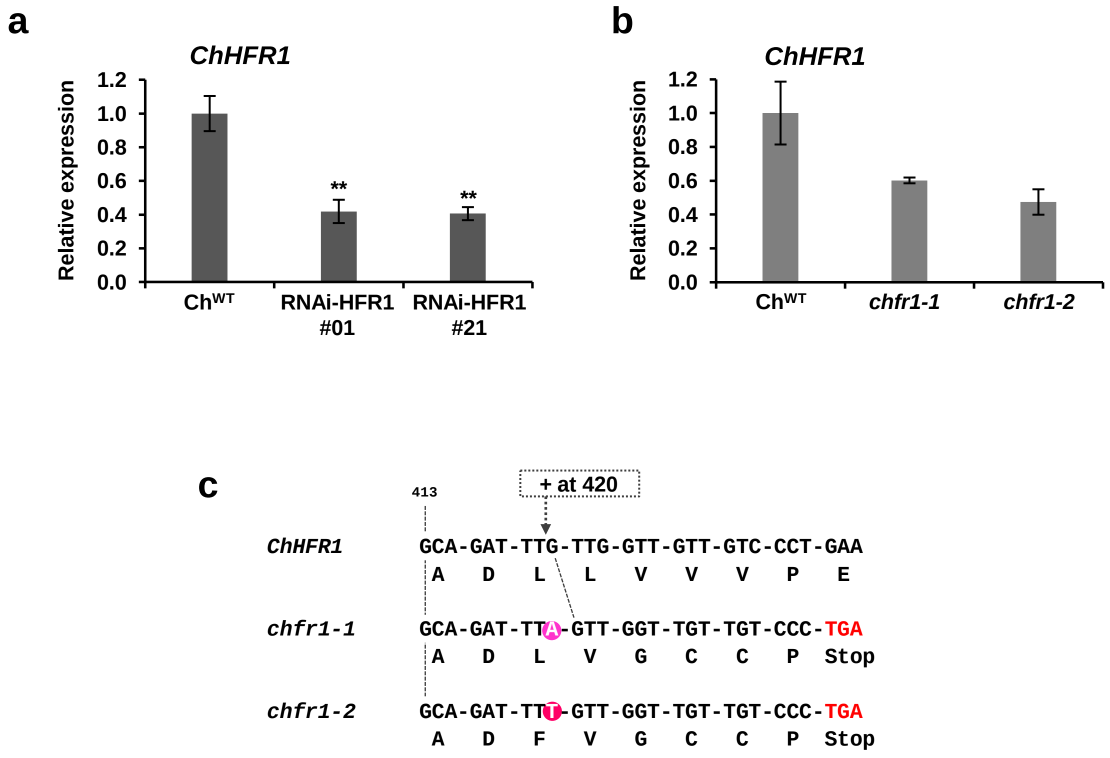


**Supplemental Figure 1. Characterization of RNAi-HFR1 and *chfr1* mutants in *C. hirsuta*.** Relative expression levels of *ChHFR1* gene, normalized to *EF1α* in Ch$^{WT}$, **(a)** two RNAi-HFR1 lines (#01 and #21) and **(b)** the two *chfr1* mutants of *C. hirsuta.* Seedlings were grown for 7 days in W. Expression values are the mean ± SE of three independent biological replicates relative to Ch$^{WT}$. **(c)** The two identified *chfr1-1* and *chfr1-2* mutants have one nucleotide insertion at position 420 of the *ChHFR1* ORF, which leads to a frame shift and a premature stop codon.

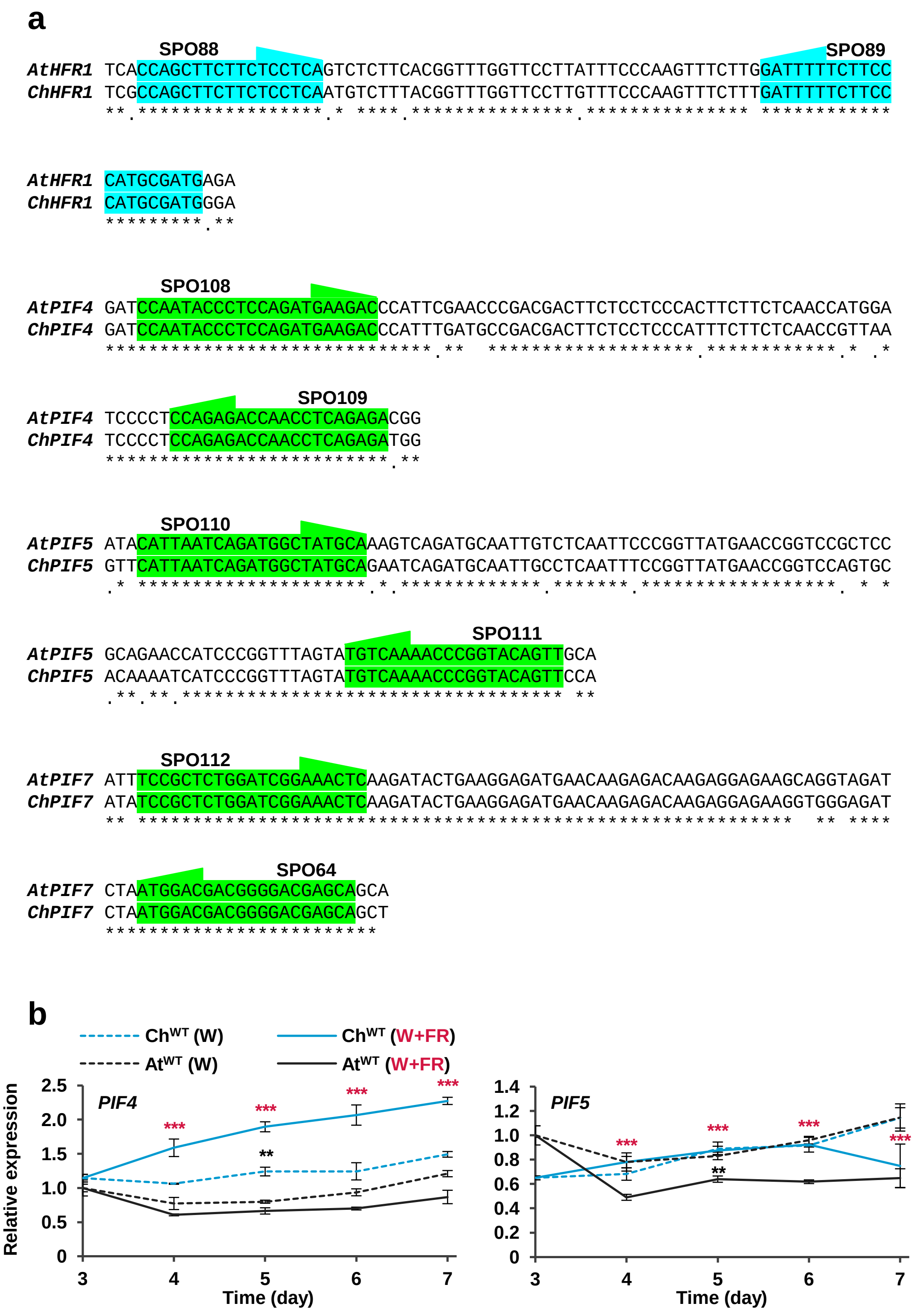


**Supplemental Figure 2. Alignments of *HFR1*, *PIF4*, *PIF5* and *PIF7* partial DNA sequences in *A. thaliana* and *C. hirsuta* (a).** Location of shared primers and amplicons used for comparison of expression levels by RT-qPCR between species. **(b)** Transcript abundance of *PIF4* and *PIF5*, normalized to *YLS8*, *SPC25* and *EF1α* in $Ch^{WT}$ and $At^{WT}$ grown as in Figure 2. Expression values are the means ± SE of three independent biological replicates relative to the data of $At^{WT}$ grown in continuous W at day 3. Asterisks mark significant differences (2-way ANOVA: * p-value <0.05, ** p-value <0.01, *** p-value <0.001) between $Ch^{WT}$ and $At^{WT}$ when grown under W (black asterisks) or W+FR (red asterisks).

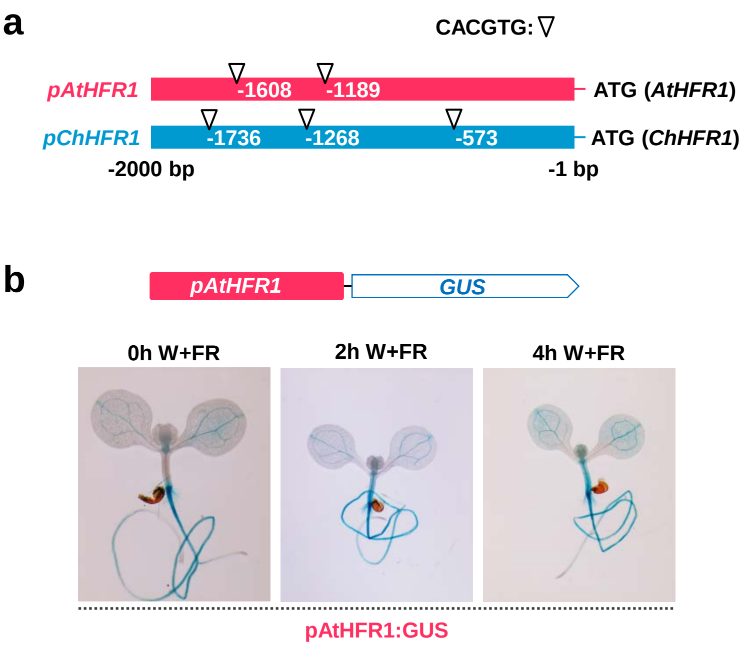


**Supplemental Figure 3. The 2 Kbp *AtHFR1* promoter is shade responsive in *A. thaliana*. (a)** Cartoon of *HFR1* promoters from *A. thaliana* (*pAtHFR1*) and *C. hirsuta* (*pChHFR1*). These promoters cover 2000 bp from the beginning of the translation start of the two *HFR1* genes. The positions of G-boxes (CACGTG) are indicated with arrows. **(b)** GUS staining of representative *A. thaliana* seedlings expressing *GUS* under the *pAtHFR1* (line #03). Seven-day-old W-grown seedlings were treated with W+FR for the indicated amount of time.

**a**

```
AtHFR1 ---------- ---------- ---------- MSNNQAFMEL GWRNDVGSLA VKDQGMMSER ARSDEDRLIN
ChHFR1 MGFPFSRTNL KSPKKNSFLK FSVPDFSLVN MFSNQDFMEL GWRNEVESLA LKDHG-ITDI ARSDEDRLIN
                                   * .** **** ****:* *** :**:* :::  **********

AtHFR1 GLKWGYGYFD HDQTDN-YLQ IVPEIHKEVE NAK-EDLLVV VPDEHSETDD HH--HIKDFS ERSDHRFYLR
ChHFR1 GLKWSYGYFG HDQTHNDHHQ IVPEIQKEER LLKTADLLVV VPDEHSETGD YHHDHIDDYS DSSDNLCYLR
       ****.****. ****.* : * *****:** .   *  ***** ********.* :*  **.*:* : **:  ***

AtHFR1 NKHENPKKRR IQVLSSDDES EEFTREVPSV TRKGS-KRRR RDEKMSNKMR KLQQLVPNCH KTDKVSVLDK
ChHFR1 NKHENPKRRR VQIW-SDEES YGFTREVPSL TRKGSKKRRR RDDELSNKMS TLQELLPNCH KADTVSVLDN
       *******:** :*:  **:**   *******: ***** **** **:::****  .**:*:**** *:*.*****:

AtHFR1 TIEYMKNLQL QLQMMSTVGV NPYFLPATLG FGMHNH-MLT AMASAHGLNP ANHMMPSPLI PALNWPLPPF
ChHFR1 AIEYMKNLQL QLQVMSAMGM NPYFPPATLD FGMSNHYMLT AMALAHIQNP AYQKTSSPLI PASNWPLLPF
       :********* ***:**::*: *********. *** ** *** *** **  ** * :  .**** ** **** **

AtHFR1 TNISFPHSSS QSLFLTTSSP ASSPQSLHGL VPYFPSFLDF SSHAMRRL
ChHFR1 TN-------- -PLFLTTASP ASSPQCLYGL VPCFPSFFDF SSHAMGRL
       **          .*****:** *****.*:** ** ****:** ***** **
```

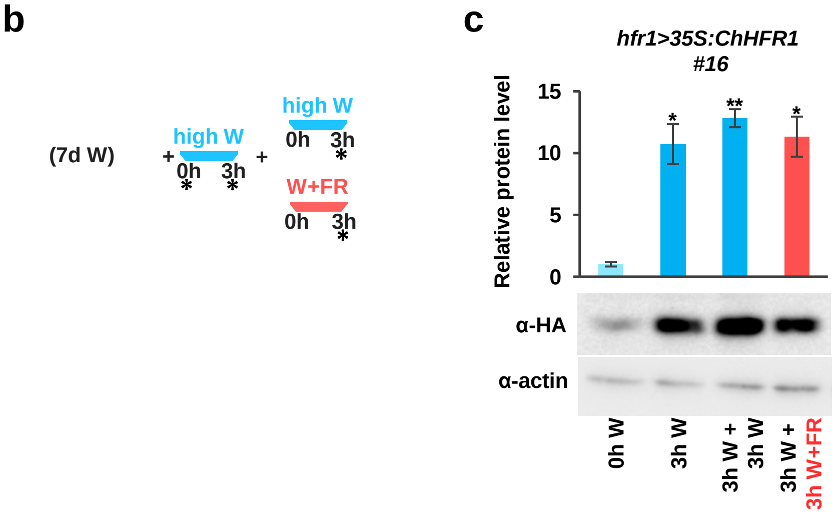


**Supplemental Figure 4. ChHFR1 protein accumulates in high W. (a)** Alignment of AtHFR1 and ChHFR1 protein sequences. Putative COP1 interacting motifs, defined in AtHFR1, are indicated in blue. **(b)** Cartoon representing the light treatments given to seedlings to estimate relative HFR1-3xHA levels. Seedlings grown for 7 d in low W (~20 µmol m$^{-2}$ s$^{-1}$, R:FR≈6.4) were first moved to high W (~100 µmol m$^{-2}$ s$^{-1}$, R:FR≈3.9) for 3 h and then either transferred to high W (control) or high W+FR (R:FR≈0.06) for 3 h. Seedling samples were collected at the time points indicated with asterisks. **(c)** Relative HFR1-3xHA protein levels of *hfr1>35S:ChHFR1* seedlings (line #16) grown as indicated in **b**, with a representative immunoblot in a lower panel. Relative protein levels are the mean ± SE of three independent biological replicates relative to the data point of 0 h in high W (0 h W). Asterisks mark significant differences in protein levels (Student *t*-test: ** p-value <0.01; * p-value <0.05) relative to the 0 h W value.

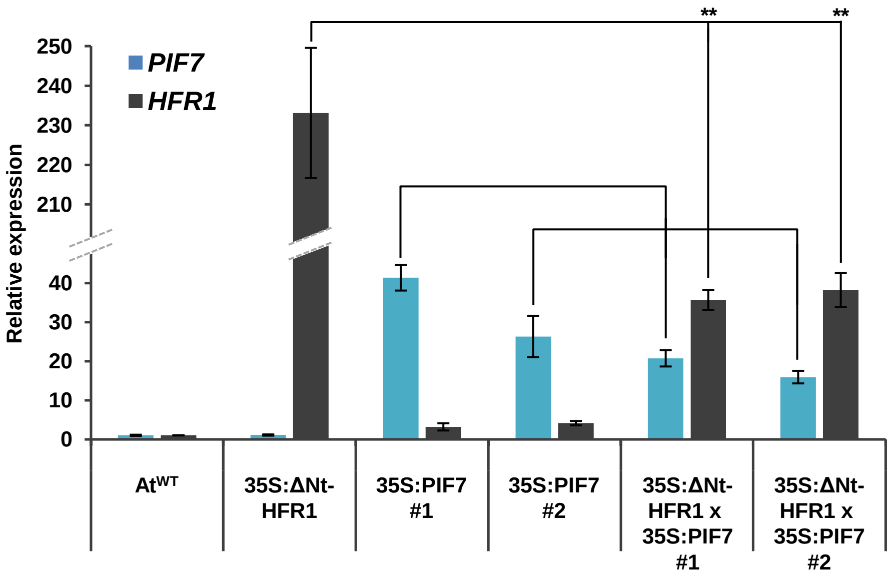


**Supplemental Figure 5. Relative expression levels of *AtHFR1* and *AtPIF7* genes in transgenic lines overexpressing *GFP-ΔNt-HFR1* and/or *PIF7-CFP*.** Relative expression, normalized to *UBQ10*, was estimated in seedlings grown for 7 days in W. Expression values are the mean ± SE of three independent biological replicates relative to At$^{WT}$. Asterisks mark significant differences in expression (Student *t*-test: ** p-value <0.01; * p-value <0.05) relative to 35S:GFP-ΔNt-HFR1-GFP or 35S:PIF7-CFP values.
